# Low-affinity ligands of the epidermal growth factor receptor are long-range signal transmitters during collective cell migration of epithelial cells

**DOI:** 10.1101/2024.09.25.614853

**Authors:** Eriko Deguchi, Shuhao Lin, Daiki Hirayama, Kimiya Matsuda, Akira Tanave, Kenta Sumiyama, Shinya Tsukiji, Tetsuhisa Otani, Mikio Furuse, Alexander Sorkin, Michiyuki Matsuda, Kenta Terai

**Author notes:** **Corresponding Author** Kenta Terai − Department of Pathology and Biology of Diseases, Graduate School of Medicine, Kyoto University, Kyoto 606-8501, Japan.

## Abstract

Epidermal growth factor receptor ligands (EGFRLs) consist of seven proteins. In stark contrast to the amassed knowledge concerning the epidermal growth factor receptors themselves, the extracellular dynamics of individual EGFRLs remain elusive. Here, employing fluorescent probes and a tool for triggering ectodomain shedding of EGFRLs, we show that EREG, a low-affinity EGFRL, exhibits the most rapid and efficient activation of EGFR in confluent epithelial cells and mouse epidermis. In Madin-Darby canine kidney (MDCK) renal epithelial cells, EGFR- and ERK-activation waves propagate during collective cell migration in an ADAM17 sheddase- and EGFRL-dependent manner. Upon induction of EGFRL shedding, radial ERK activation waves were observed in the surrounding receiver cells. Notably, the low-affinity ligands EREG and AREG mediated faster and broader ERK waves than the high-affinity ligands. The integrity of tight/adherens junctions was essential for the propagation of ERK activation, implying that the tight intercellular spaces prefer the low-affinity EGFRL to the high-affinity ligands for efficient signal transmission. To validate this observation *in vivo*, we generated EREG-deficient mice expressing the ERK biosensor and found that ERK wave propagation and cell migration were impaired during skin wound repair. In conclusion, we have quantitatively demonstrated the distinctions among EGFRLs in shedding, diffusion, and target cell activation in physiological contexts. Our findings underscore the pivotal role of low-affinity EGFRLs in rapid intercellular signal transmission.

## Introduction

The epidermal growth factor receptor (EGFR)-Ras-Extracellular signal-regulated kinase (ERK) signaling pathway governs a plethora of biological phenomena, encompassing cell proliferation, differentiation, and tumorigenesis (Linggi and Carpenter, 2006; Nishida and Gotoh, 1993; Yarden and Sliwkowski, 2001). The EGFR ligands (EGFRLs) consist of seven proteins and are categorized into two groups based on their receptor-binding affinity (Harris et al., 2003). The high-affinity ligands, with apparent Kd values ranging from 0.1 to 1 nM, are epidermal growth factor (EGF), transforming growth factor-α (TGFα), betacellulin (BTC), and heparin-binding EGF-like growth factor (HBEGF). In contrast, epiregulin (EREG), epigen (EPGN), and amphiregulin (AREG) are the low-affinity ligands, exhibiting affinities 10- to 100-fold lower than their high-affinity counterparts (Jones et al., 1999). Recent investigations have illuminated that the receptor-binding affinity differentially stabilizes EGFR dimers and thereby plays a pivotal role in eliciting distinct cellular responses (Freed et al., 2017; Hu et al., 2022). EGFRLs can also be categorized based on their sensitivity to sheddases (Sahin et al., 2004), their bioactivity in promoting cell growth and migration (Singh et al., 2016; Wilson et al., 2009), and their endocytic sorting (Roepstorff et al., 2009).

In addition to traditional fluorescently tagged EGFRLs, advanced probes have been developed to probe the characteristics of EGFRLs. First, the sensitivity to sheddases was examined by fusing peptide tags or alkaline phosphatase to the extracellular domain of EGFRLs (Dang et al., 2013; Hinkle et al., 2004; Sahin et al., 2004; Tokumaru et al., 2000). Studies using these probes demonstrated that HBEGF, TGFα, EREG, AREG, and EPGN, but not EGF or BTC, undergo shedding by a disintegrin and metalloprotease 17 (ADAM17). Secondly, fluorescent proteins have been fused to the C-terminus of EGFRLs to investigate the intracellular trafficking of pro-EGFRLs (Gephart et al., 2011; Prince et al., 2010; Singh et al., 2015; Singh et al., 2013). Thirdly, several groups have created probes by attaching fluorescent proteins to the extracellular domain of proEGFRLs {Inoue, 2013 #19;Kamezaki, 2016 #20;Bunker, 2021 #21}. While these probes were employed to monitor the cleavage efficiency by ADAM17, they did not provide insights into the extracellular dynamics of the shed EGFRL.

Notably, a substantial portion of current knowledge about the biological effects of individual EGFRLs is derived from experiments involving the bath application of recombinant EGFRLs to cultured cells. This approach, however, leaves unresolved questions concerning shedding, diffusion, and target cell activation within physiological contexts. A unique model system for investigating the physiological roles of EGFRLs is the collective cell migration exhibited by Madin-Darby canine kidney (MDCK) cells (Hirashima et al., 2023), wherein recurrent waves of ERK activation propagate from the leader cells to the follower cells (Aoki et al., 2017; Aoki et al., 2013; Hiratsuka et al., 2015). The propagation of these ERK activation waves, or simply ERK waves hereafter, has been shown to hinge upon ADAM17-mediated shedding of EGFRLs and intercellular mechanochemical force (Hino et al., 2022; Hino et al., 2020). Later experiments demonstrated that all four EGFRLs expressed in MDCK cells—namely EGF, HBEGF, TGFα, and EREG—collectively contribute to the ERK activation waves (Lin et al., 2022). It has also been elucidated that MDCK cells express EGFR, ErbB2, and ErbB3, but not ErbB4, with EGFR assuming the principal role in the propagation of ERK waves (Matsuda et al., 2023). Nevertheless, comprehensive analyses of individual EGFRLs have not been performed due to the absence of suitable probes for tracking EGFRLs and methods for inducing EGFRL shedding.

Here, we report our design of a series of EGFRL probes, named EGFRL-ScNeos, for visualization of the shedding and extracellular dynamics of individual EGFRLs. By using these probes in tandem with a chemical biology tool for eliciting EGFRL shedding, we elucidate distinctive characteristics inherent to each EGFRL. Remarkably, our observations reveal that EREG, one of the low-affinity EGFRLs, unexpectedly functions as a long-range signaling mediator, traversing the intercellular milieu beneath the tight/adherens junctions.

## Results

### EGFRL-ScNeos, dual color probes, visualize the shedding of EGFRLs and stimulate EGFR

We developed a series of seven dual-color fluorescent probes, EGFRL-ScNeos, to visualize the dynamics of the seven EGFRLs in live cells (Fig. 1A, 1B). Each EGFRL-ScNeo included the red fluorescent protein mScarlet and the green fluorescent protein mNeonGreen at the extracellular and cytoplasmic domains, respectively. mScarlet was inserted at the N-terminus of the EGF domain. In the case of the probes for EGF, EREG, and TGFα, mScarlet was sandwiched with linker peptides. mNeonGreen was fused to the C-terminus of EGFRL except in the case of the TGFα probe, for which mNeonGreen was inserted before the C-terminal tandem valine residues required for intracellular trafficking (Briley et al., 1997; Dempsey et al., 2003). A probe for NRG1, a ligand for ErbB3 and ErbB4, was also developed according to the method in a previous paper (Kamezaki et al., 2016). As the control protein that is not cleaved by ADAM proteases, we employed the transmembrane protein Necl-5/PVR/CD155, simply Necl-5 hereafter (Ichise et al., 2022).

**Figure 1.**
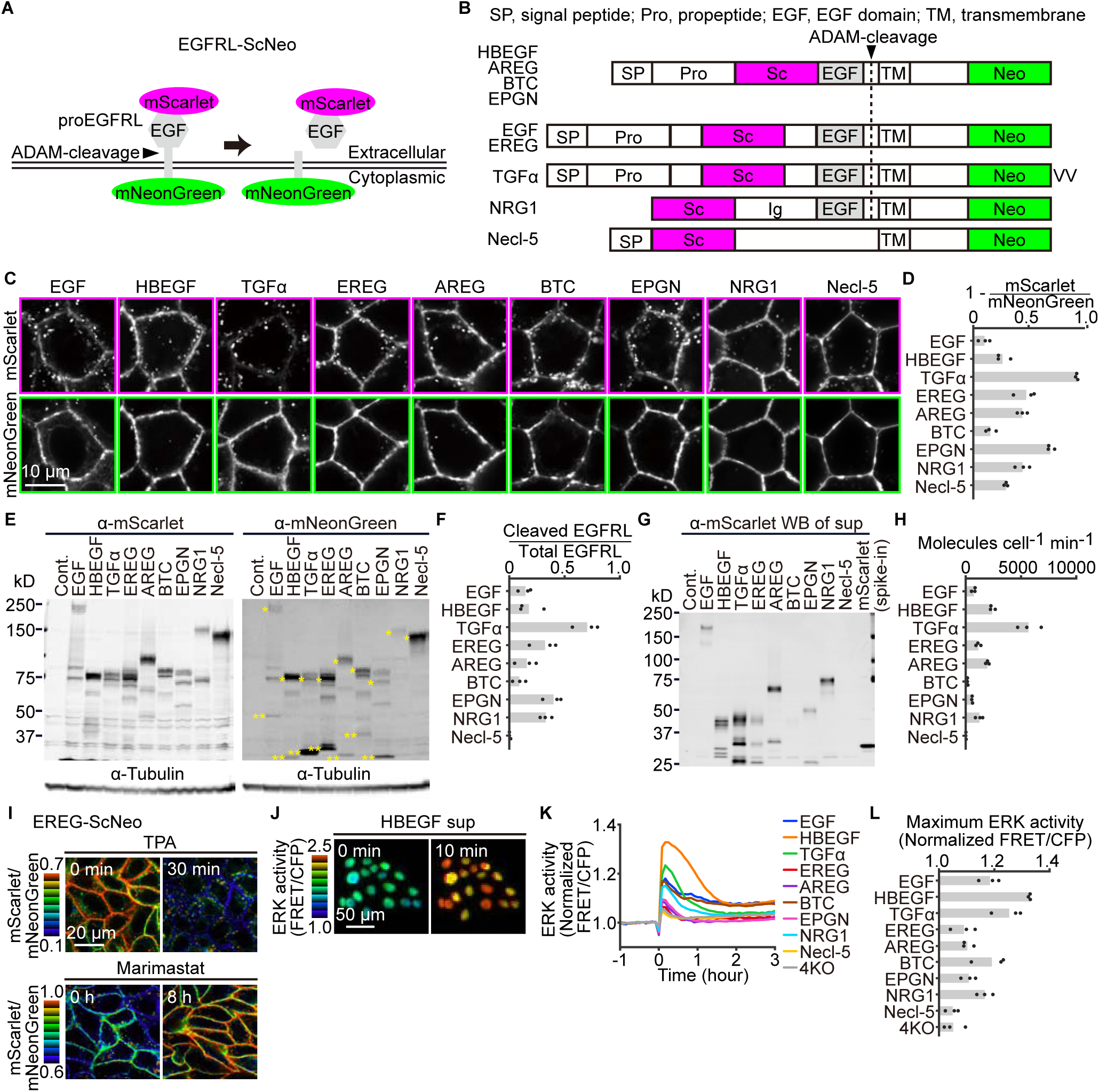
EGFRL-ScNeos, dual color probes, visualize shedding of EGFRLs and stimulate EGFR. **(A)** Each EGFRL-ScNeo included the mScarlet red fluorescent protein and the mNeonGreen green fluorescent protein in the extracellular and cytoplasmic domains, respectively. ADAM-family proteases shed mScarlet-fused EGFRL. **(B)** mScarlet was inserted in front of the EGF domain. mNeonGreen was fused to the C terminus except in the case of TGFα, in which mNeonGreen was inserted before the C-terminal tandem valine residues. Note that the pro-NRG1 lacks the signal peptide and that Necl-5 lacks the ADAM cleavage site. SP, signal peptide; Pro, propeptide; EGF, EGF domain; TM, transmembrane domain; Ig, immunoglobulin-like domain. **(C)** XY confocal images of EGFRL-ScNeos expressed in MDCK cells. **(D)** mScarlet/mNeonGreen fluorescence ratio of the cell membrane in (C). The bar graphs show the mean values. Each dot represents the average value for one experiment. n > 100 cells for each condition. **(E)** Western blotting of total cell lysates of EGFRL-ScNeo-expressing cells. Membranes were probed with an anti-mScarlet or an anti-mNeonGreen antibody. *Full-length EGFRL-ScNeo; **Cytoplasmic domain fused with mNeonGreen. **(F)** The proportion of cleaved EGFRL-ScNeo in (E). The intensity of cytoplasmic domain fused with mNeonGreen (**) was divided by the intensity of the total detected bands for each ligand. The bar graphs show the mean values. Each dot indicates an independent experiment. **(G)** Western blotting of supernatants of EGFRL-ScNeo-expressing cells. Membranes were probed with an anti-mScarlet antibody. **(H)** The production rates of EGFRL from a single EGFRL-ScNeo-expressing cell were calculated from the image shown in (G). The bar graphs show the mean values. Each dot indicates an independent experiment. **(I)** Representative mScarlet/mNeonGreen ratio images of EREG-ScNeo-expressing MDCK cells upon treatment with 10 nM TPA (Video S1) or 10 μM Marimastat (Video S2). **(J)** Representative images of ERK activity (FRET/CFP) in MDCK-4KO cells expressing EKARrEV-NLS stimulated with the supernatant of MDCK-4KO cells expressing HBEGF-ScNeo. **(K)** MDCK-4KO-EKARrEV-NLS cells observed under a fluorescent microscope were stimulated with the supernatant of MDCK-4KO cells expressing EGFRL-ScNeo. The FRET/CFP ratio was normalized to the average values over the 60 min-observation period before stimulation. ERK activity (normalized FRET/CFP) was quantified and plotted as a function of time. Data were pooled from two independent experiments. n > 1000 cells for each condition. **(L)** Maximum ERK activity (normalized FRET/CFP) in the time course of average ERK activity in (K). The bar graphs show the mean values. Each dot represents the average value for one experiment. n > 1000 cells from three independent experiments were used for each condition.

We first evaluated the subcellular localization of EGFRL-ScNeos in confluent MDCK cells (Fig. 1C). The cytoplasmic mNeonGreen signal was restricted to the plasma membrane, whereas the extracellular mScarlet signal was observed both in the plasma membrane and endosomes, suggesting that the cleaved extracellular domains were engulfed and sorted to the endosomes by the EGFRL-ScNeo-producer cells themselves or neighboring cells. In the XZ section, all probes were localized primarily at the basolateral plasma membrane and to a lesser extent at the apical membrane (Fig. S1A). The relative efficiency of cleavage was quantified by measuring the fluorescence intensity of extracellular mScarlet and cytoplasmic mNeonGreen on the plasma membrane (Fig. 1D). In the Western blot analysis of cell lysates (Fig. 1E), the uncleaved pro-EGFRLs were detected both in anti-mScarlet and anti-mNeonGreen blots. The bands expected from molecular weights are marked with single asterisks on the left.

Meanwhile, the cleaved cytoplasmic domain of EGFRLs were detected in anti-mNeonGreen, but not anti-mScarlet blot. The bands are marked with double asterisks on the left. We assumed that the other minor bands are generated by incomplete cleavage of multiple protease sensitive sites or glycosylation (Hinkle et al., 2004; Kochupurakkal et al., 2005; Le Gall et al., 2003). We estimated the fraction of cleaved probes based on the Western blot with anti-mNeonGreen antibody (Fig. 1F). Secretion of the cleaved EGFRL into the medium was also examined by quantifying EGFRL in the supernatant (Fig. 1G). With these data in hands, we calculated the production rate of EGFRL secreted into the medium (Fig. 1H). In conclusion, among the seven EGFRLs TGFα was most efficiently cleaved and secreted into the medium in the absence of any stimulation. Of note, the subcellular localization of the probes was not significantly affected when cells were grown on permeable supports (Fig. S1B).

Next, the ADAM17-dependent shedding of EGFRL-ScNeos was quantified by the plasma membrane mScarlet/mNeonGreen ratio before and after the addition of a stimulator, 12-O-tetradecanoylphorbol 13-acetate (TPA), or an inhibitor, Marimastat (Fig. 1I, S1C-D, Video S1, S2). The mScarlet/mNeonGreen ratios of EREG, AREG, and EPGN, but not EGF, HBEGF, TGFα, BTC, and NRG1, were significantly decreased by TPA. The mScarlet/mNeonGreen ratios of all EGFRLs except for EGF and BTC were significantly increased by Marimastat. This observation is consistent with the previous reports that, except for EGF and BTC, EGFRLs are sensitive to ADAM17 (Sahin et al., 2004). Although TPA is known to induce HBEGF shedding with high efficiency (Goishi et al., 1995), mScarlet/mNeonGreen ratio of HBEGF was not decreased by TPA. We can offer two possible explanations. First, as suggested in a previous report (Dang et al., 2013), TPA may alter the substrate specificity of ADAM17. Secondly, the heparin-binding nature of HBEGF could have prevented the cleaved HBEGF from diffusing, thereby preventing the decrease in mScarlet fluorescence.

Finally, we investigated whether the cleaved EGFRL-ScNeos retained biological activity to stimulate EGFR. For this purpose, each EGFRL-ScNeo was stably expressed in MDCK-4KO cells in which all four EGFRL genes expressed in MDCK cells, EGF, HBEGF, TGFα, and EREG, were knocked out (Lin et al., 2022). The culture supernatants of these EGFRL-ScNeo-expressing cells were applied to MDCK cells expressing the FRET biosensor for ERK (Fig. 1J-L). All supernatants derived from the EGFRL-ScNeo-expressing cells transiently stimulated ERK (Fig. 1J), although the level of maximum activation was different for each EGFRL (Fig. 1K). In this assay, we observed that HBEGF stimulated ERK most efficiently. We obtained similar results by using commercially available recombinant EGFRLs, indicating that the mScarlet-tagged EGFRLs retained the biological activity (Fig. 1L, Fig. S1E). This observation was also confirmed by immunoblotting with the anti-phospho-EGFR antibody (Fig. S1F). Thus, we concluded that EGFRL-ScNeos retain its biological activity and reflect the dynamics of EGFRLs.

### EGFRL-ScNeo highlights short- and long-range EGFRLs

Using the above-described EGFRL-ScNeos, we addressed the question of how far each EGFRL travels toward the surrounding cells after being shed from the producer cells. The EGFRL-ScNeo-producer MDCK cells were co-cultured with an excess of wild-type MDCK receiver cells (Fig. 2A). The mScarlet signal was used to track each EGFRL (Fig. 2B-D). As expected, the mScarlet signal was not observed in the cells surrounding the Necl5-ScNeo-expressing cells, whereas mScarlet signals were detected up to 80 μm from the HBEGF-ScNeo-expressing producer cells. To a lesser extent, mScarlet signals were detected around cells expressing NRG1-ScNeo, EGF-ScNeo, and AREG-ScNeo. The mScarlet signals were fainter in cells surrounding the producer cells of TGFα, EREG, and BTC, probably reflecting the low affinity to the EGFR or low cleavage rate of these EGFRLs. The mScarlet signals were below the quantifiable level in cells surrounding the EPGN-producer cells, probably reflecting the low affinity to the EGFR. The mScarlet signals in the receiver cells were abolished when ADAM17 was eliminated from the producer cells, confirming that the signals were derived from the ADAM17-cleaved EGFRL-ScNeo (Fig. S2).

**Figure 2.**
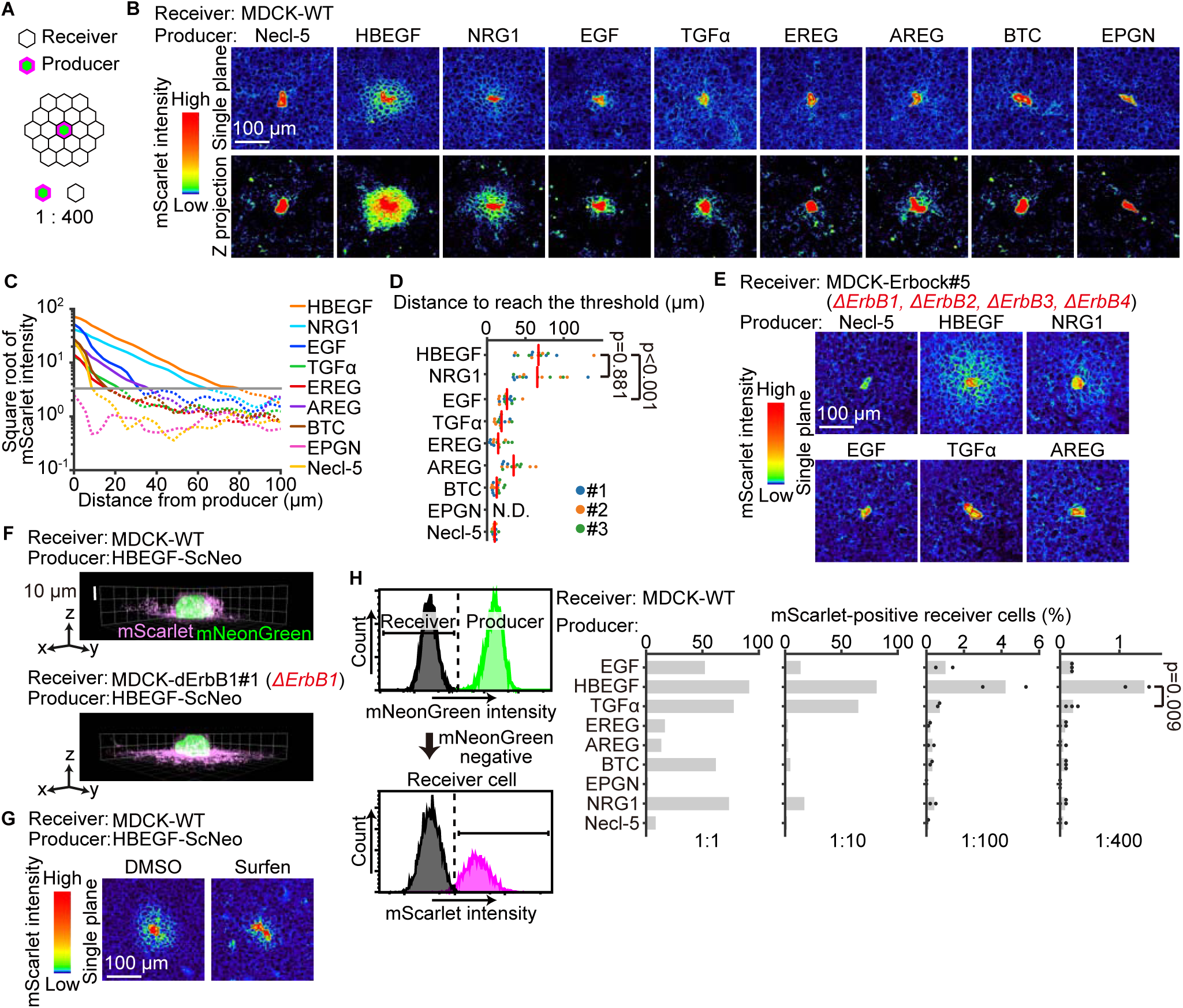
EGFRL-ScNeo highlights short and long-range EGFRLs. **(A)** Schematic of the co-culture experiment. EGFRL-ScNeo-expressing cells and parental MDCK cells were used as the producer and receiver cells, respectively. The producer and receiver cells were co-cultured at a ratio of 1:400. **(B)** From mScarlet confocal images of cells, a single representative plane and a Z projection image of 20 slices are shown for each EGFRL. The central signals above the threshold are from the producer cells. **(C)** mScarlet signal gradient from the producer cells in (B). The square root of mean values of mScarlet intensity in the receiver cells was plotted over the distance from the producer cells. The data were normalized to the mNeonGreen intensity of producer cells. The gray bar represents the threshold of detectable mScarlet signals. The data were pooled from three independent experiments. Each experiment includes 5 images. **(D)** Distance from producer cells to reach the threshold indicated in (C). Approximate curves were set from the first five points in the mScarlet decay curve in (C). The distance to reach the threshold was then calculated. The red bars represent the means. n = 5 images from three independent experiments, depicted by the three colors. **(E)** Identical to (B) except that the receiver cells were MDCK-Erbock cells, which lack all four ErbB-family receptors. **(F)** 3D images of HBEGF-ScNeo cells co-cultured with WT or ErbB1-deficient MDCK cells. **(G)** mScarlet confocal images of HBEGF-ScNeo cells co-cultured with WT MDCK cells. 3 hours after seeding, cells were supplemented with DMSO 0.1% or 5 μM surfen, and maintained for one day under the condition. **(H)** (Left) Cells prepared as in (A) were analyzed by flow cytometry analysis. The proportion of mScarlet-positive cells among mNeonGreen-negative receiver cells was quantified. (Right) The proportion of mScarlet-positive receiver cells at different producer vs. receiver ratios. EGFRL-ScNeo-expressing cells were used as producers and parental wild-type cells were used as receivers. The bar graphs show the mean values (n = 1 for 1:1 and 1:10, n = 2 for 1:100, n = 3 for 1:400). Each dot represents an independent experiment. Statistical significance was determined by unpaired two-tailed Welch’s t-test.

We then examined whether the range of the distribution depended on binding to EGFR by changing the receiver cells to Erbock cells, in which all four ErbB-family receptors are knocked out (Matsuda et al., 2023) (Fig. 2E). At first look, it appeared that the mScarlet signals were not changed significantly when HBEGF-ScNeo and NRG1-ScNeo were used. However, in looking at the tangential images, we found that HBEGF was accumulated at the basal surface of EGFR deficient cells (Fig. 2F), suggesting that HBEGF was captured by heparan sulfate proteoglycans (HSPGs) on the cell surface of the parent MDCK cells. In accordance with this hypothesis, surfen, which prevents growth factors from binding to HSPGs, significantly suppressed the mScarlet signal in the receiver cells surrounding the HBEGF producer cell (Fig. 2G).

Next, cellular uptake of EGFRL by the receiver cells was quantified by flow cytometry (Fig. 2H). When producer and receiver cells were co-cultured at a ratio of 1:1, more than 50% of the receivers were scored as mScarlet-positive with the probes for EGF, HBEGF, TGFα, BTC, and NRG1, which are classified as high-affinity ligands. At a co-culture ratio of 1:10, more than 50% of HBEGF- and TGFα-receiver cells were still mScarlet-positive, but this percentage decreased markedly at co-culture ratios of 1:100 and 1:400; only receiver cells of the HBEGF probe had a significant mScarlet-positive population under these ratios. This result implies that TGFα was diluted to below the detection limit, whereas HBEGF reached a limited number of cells over a short distance and thus retained a strong signal.

### Low-affinity EGFRLs diffuse faster and farther than high-affinity EGFRLs

The above experiments investigated the distribution of EGFRLs in the absence of stimulation, i.e., constitutive cleavage by the basal sheddase activity. Here, we examined the EGFRL dynamics upon acute ADAM17 activation by SLL-induced protein translocation (SLIPT). Briefly, bath application of a localized ligand, m^D^cTMP, anchors cytoplasmically expressed modified *Escherichia coli* dihydrofolate reductase eDHFR fused to cRaf to the inner leaflet of the plasma membrane (Suzuki et al., 2022), thereby activating ADAM17. To validate this system, we introduced eDHFR-cRaf into cells expressing TSen, a FRET probe for ADAM17 (Chapnick et al., 2015) (Fig. 3A). As anticipated, upon addition of m^D^cTMP, ADAM17 was activated in a dose-dependent manner (Fig. 3B). We then expressed AREG-ScNeo and miRFP703-eDHFR-cRaf and observed membrane translocation of miRFP703-eDHFR-cRaf and a decrease in the mScarlet/mNeonGreen ratio upon m^D^cTMP addition (Fig. 3C, Video S3), indicating that acute ADAM17 activation caused AREG shedding.

**Figure 3.**
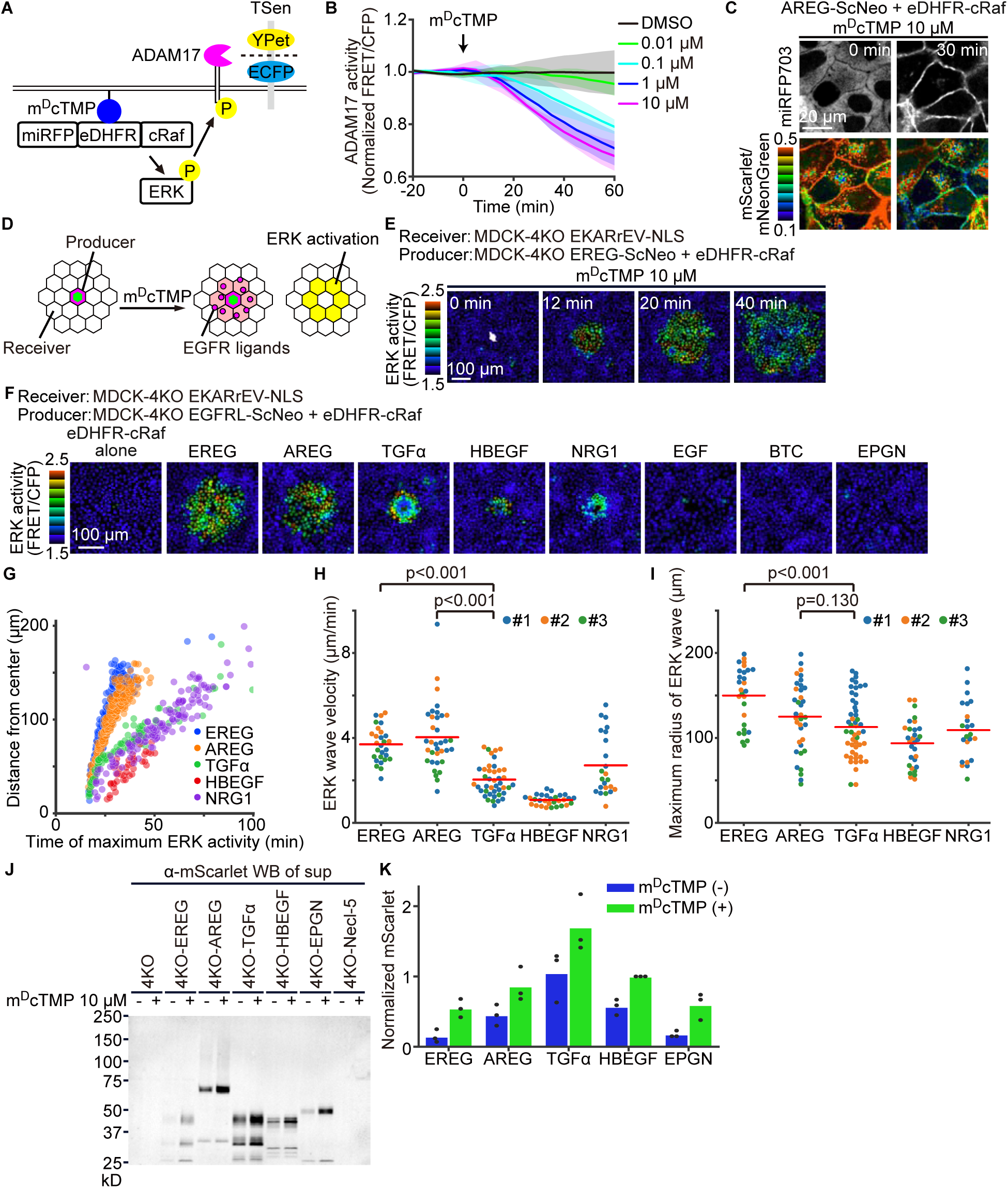
Low-affinity EGFRLs propagate ERK activation more efficiently than high-affinity EGFRLs. **(A)** Schematic of the SLL-induced protein translocation (SLIPT) system. Bath application of m^D^cTMP, which localizes at the inner leaflet of the plasma membrane and binds to *Escherichia coli* dihydrofolate reductase (eDHFR), recruits the miRFP703-eDHFR-cRaf fusion protein to the plasma membrane. cRaf at the plasma membrane stimulates ERK, which in turn activates ADAM17. The activated ADAM17 at the plasma membrane induces ectodomain shedding of EGFRL-ScNeo. The miRFP703-eDHFR-cRaf fusion protein was expressed in MDCK-4KO cells (*ΔEGF, ΔHBEGF, ΔTGFA, ΔEREG*) to generate eDHFR-cRaf cells. **(B)** TSen, a FRET biosensor for ADAM17, was introduced into the eDHFR-cRaf cells. Cells observed under a fluorescent microscope were stimulated with various concentrations of m^D^cTMP. The FRET/CFP ratio was normalized to the average values over the 20 min-observation period before stimulation. The average ERK activity (normalized FRET/CFP) was quantified and plotted as a function of time with the s.d. Data were pooled from three independent experiments. n > 100 cells for each condition. **(C)** The AREG-ScNeo probe was introduced into the eDHFR-cRaf cells. Cells were treated with 10 μM m^D^cTMP. Representative images of miRFP703 and mScarlet/mNeonGreen ratio are shown (Video S3). **(D)** EGFRL-ScNeo probes were introduced into the MDCK-4KO cells expressing eDHFR-cRaf to generate the producer cells, which were co-cultured with an excess of MDCK-4KO cells expressing a FRET biosensor for ERK, EKARrEV-NLS. **(E)** Representative time-lapse ERK activity (FRET/CFP) images. The EREG-producer cells are located at the center of the image. Surrounding cells are MDCK-4KO cells expressing EKARrEV-NLS. **(F)** Each ligand producer is located at the center of the image. Images are snapshots of Video S4 acquired 20 min after m^D^cTMP addition. **(G)** In each cell shown in (F), the time of the highest ERK activity (FRET/CFP) is plotted against the distance to the producer cells. **(H)** From the scatter plot in (G), velocities of ERK waves were calculated. Each dot indicates a single producer cell population. The red bars represent the means. n = 28 (EREG), 36 (AREG), 50 (TGFα), 30 (HBEGF), and 23 (NRG1) producer cell populations from three independent experiments, depicted by the three colors. **(I)** Maximum radius of the ERK-activated area upon EGFRL shedding. Data in (H) are used for the analysis. The red bars represent the means. Statistical significance**(J)** Analysis of the m^D^cTMP-dependent EGFRL shedding. EGFRL-ScNeo-expressing 4KO-eDHFR-cRaf cells were incubated with or without 10 μM m^D^cTMP for 1 hour. The supernatant of the cells was corrected and calculated the amount of EGFRL with an anti-mScarlet antibody. **(K)** The production rates of EGFRL. mScarlet intensities of HBEGF sup with m^D^cTMP were set as one. The bar graphs show the mean values. Each dot indicates an independent experiment.

Next, by using this system, we visualized how shed EGFRL activates EGFR in the surrounding receiver cells by measuring ERK activity using the EKARrEV FRET biosensor. Specifically, MDCK-4KO cells expressing both EGFRL-ScNeo and eDHFR-cRaf were co-cultured with EKARrEV-NLS-expressing MDCK-4KO receiver cells (Fig. 3D). Note that the effect of endogenous EGFRLs was eliminated in this assay because both producer and receiver cells lacked endogenous EGFRLs. Upon addition of m^D^cTMP, ERK activation was initiated around the producer cells, followed by the radial spread of ERK activation when the producer cells expressed EREG, AREG, TGFα, HBEGF, and NRG1, but not when they expressed EGF, BTC, and EPGN (Fig. 3E, 3F, Video S4). The lack of response of EPGN was likely due to the low level of the uncleaved form before stimulation (Fig. S1C). The lack of response of EGF and BTC agreed with the insensitivity to ADAM17 inhibitor (Fig. S1D). The essential role of ADAM17 in the producer, but not the receiver cells was confirmed by using ADAM17-deficient MDCK cells (Fig. S2). The ERK wave velocities of the low-affinity ligands EREG and AREG, ∼4 μm min^-1^, were approximately four times higher than those of the high-affinity ligands HBEGF and TGFα, 1 to 2 μm min^-1^ (Fig. 3G, 3H). The area of ERK-activated cells was also larger for the low-affinity ligands than for the high-affinity ligands, though not significantly larger in the case of AREG (Fig. 3I). Note that the amount of EGFRLs secreted from the low-affinity ligand-expressing producer cells was at a similar level to those from the high-affinity ligand-expressing producer cells (Fig. 3J and 3K). This observation suggests that in the context of the confluent epithelial cell layer, the low-affinity EGFRLs transmit signals to the distant cells faster than the high-affinity EGFRLs.

In EGFR-deficient cells, the ERK activation wave was abolished when the producer cells expressed AREG, TGFα, or HBEGF, partially suppressed when the producer cells expressed EREG, and not affected when the producer cells expressed NRG1 (Fig. S3A). In the latter two cell lines, further deletion of ErbB3 and ErbB4 eliminated the ERK waves (Fig. S3B). These observations are consistent with the previous reports showing the binding of EREG and NRG1 to ErbB3 and ErbB4(Wilson et al., 2009). Deletion of ADAM17 in the producer cells abolished the generation of ERK waves from all producer cell lines (Fig. S3C), confirming that the SLIPT system triggers EGFRL shedding via ADAM17. To exclude the possibility that the fusion of mScarlet to EGFRL affects the binding properties, we designed probes without extracellular mScarlet (Fig. S3D). We found no significant difference in the velocity of ERK waves between the probes with and without extracellular mScarlet (Fig. S3E). Because cell–cell traction force contributes to ERK waves during collective cell migration(Hino et al., 2020), we perturbed actin polymerization but did not find any effect on m^D^cTMP-stimulated ERK waves (Fig. S3F), indicating that the propagation of ERK activation was primarily mediated by the diffusion of EGFRLs under the present condition.

### The diffusion of EGFRLs in the intercellular space is regulated by the affinity to and the density of EGFR on the basolateral plasma membrane

Why do the ERK waves elicited by the low-affinity EGFRLs propagate faster than those elicited by the high-affinity EGFRLs? Among the high-affinity EGFRLs, TGFα is unique in that it is sorted to the basolateral membrane in a Naked2-dependent manner (Li et al., 2004). To exclude the possibility that the cytoplasmic domain affects the velocity of ERK waves, we used a TGFα-EREG chimera that carries the extracellular domain of TGFα and the cytoplasmic domain of EREG (Fig. 4A). We found that the velocity of the ERK wave induced by the TGFα-EREG chimera was as slow as that induced by the authentic TGFα, supporting the notion that the high affinity to EGFR underlies the slow wave propagation (Fig. 4B). If the high affinity to EGFR slows down the wave propagation, the density of EGFR on the plasma membrane may also affect the ERK wave velocity. Indeed, overexpression of human EGFR in MDCK cells slowed down the EREG-induced ERK wave to the level of the HBEGF-induced ERK wave (Fig. 4C). Thus, in the tight intercellular spaces, EGFRLs are rapidly sequestered depending on the affinity to and the density of EGFR.

**Figure 4.**
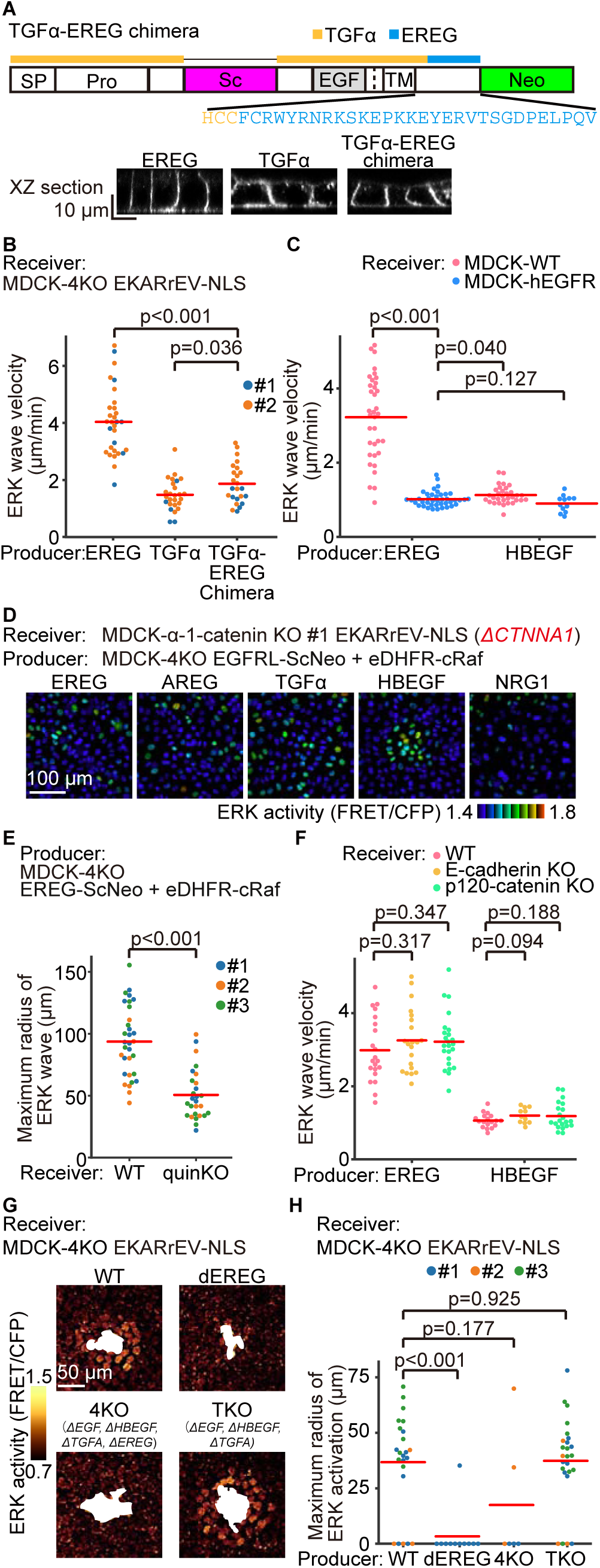
Diffusion of EGFRL in the intercellular space is regulated by the affinity to and the density of EGFR on the basolateral plasma membrane. **(A)** (Upper) A schematic of the TGFα-EREG chimera of extracellular TGFα and cytoplasmic EREG. (Bottom) mNeonGreen XZ images of EREG, TGFα, and a TGFα-EREG chimera. **(B)** The velocity of ERK wave propagated from EREG-, TGFα-, or a TGFα-EREG chimera-producer cells to MDCK-4KO-EKARrEV-NLS receiver cells. Each dot indicates a single producer cell population. The red bars represent the means. N = 30 (EREG), 26 (TGFα), and 24 (TGFα-EREG chimera) producer cell populations from two independent experiments, depicted by the two colors. **(C)** The velocity of ERK wave propagated from EREG- or HBEGF-producer cells to WT or EGFR-overexpressing receiver cells. Each dot indicates a single producer cell population. The red bars represent the means. n = 35 (EREG, WT), 41 (EREG, EGFR O/E), 32 (HBEGF, WT), and 12 (HBEGF, EGFR O/E) producer cell populations from two independent experiments. **(D)** Representative images of ERK activity (FRET/CFP) in MDCK-α-1-catenin KO receiver cells. Each ligand producer cell is located in the center of the image. Images were acquired 30 min after m^D^cTMP addition (Video S5). **(E)** Maximum radius of ERK wave propagated from EREG-producer cells to MDCK-II-WT or MDCK-II-quinKO receiver cells. Each dot indicates a single producer cell population. The red bars represent the means. n = 38 (WT) and 42 (quinKO) producer cell populations from three independent experiments, depicted by the three colors. **(F)** The velocity of ERK wave propagated from EREG- or HBEGF-producer cells to WT, E-cadherin KO, and p120-catenin KO receiver cells. Each dot indicates a single producer cell population. The red bars represent the means. n = 21 (EREG, WT), 21 (EREG, E-cadherin KO), 24 (EREG, p120-catenin KO), 18 (HBEGF, WT), 11 (HBEGF, E-cadherin KO), and 21 (HBEGF, p120-catenin KO) producer cell populations from two independent experiments. **(G)** Representative images of ERK activity (FRET/CFP) in MDCK-4KO-EKARrEV-NLS receiver cells. Each ligand producer cell is located in the white area of the image. The producer cells expressing eDHFR-cRaf are derived from the cell lines listed at the top of each image. Images were acquired 20 min after m^D^cTMP addition. **(H)** Maximum radius of ERK wave propagation in (G). Each dot indicates a single producer cell population. The red bars represent the means. n = 23 (WT), 25 (TKO) producer cell populations from three independent experiments. n = 6 (4KO) producer cell populations from two independent experiments. n = 11 (dEREG) producer cell populations from a single experiment. Statistical significance was determined by unpaired two-tailed Welch’s t-test.

What happens when the barrier of the intercellular space is perturbed? When adherens junctions were abolished by knocking out of α-1-catenin (*CTNNA1*), the ERK wave propagation was abolished except for HBEGF (Fig. 4D, Video S5). We previously demonstrated that the deletion of α-1-catenin did not influence the sensitivity to growth factors (Hino et al., 2020). Thus, except for HBEGF, which is sequestered by HSPGs, the adherens junction is essential for the ERK wave. When the tight junction was abolished in claudin quinKO MDCK II cells established by knockout of claudin-1, -2, -3, -4, and -7 genes (Otani et al., 2019), the wave propagation by EREG was significantly, but not completely, abolished (Fig. 4E). Similar experiments were performed by using E-cadherin-knockout cells and p120 catenin-knockout cells without any significant difference; thus, E-cadherin and p120 catenin are not essential for maintaining EREG within the intercellular compartments (Fig. 4F). Apico-basal polarity and tight junction formation as examined by immunostaining of ZO-1 and gp135 was lost only in *CTNNA1*-deficient cells (Fig. S4), suggesting that in addition to the presence of tight junction tight sealing by claudin proteins is required for the signal propagation by EREG.

To confirm that the rapid propagation of ERK waves is primarily mediated by EREG in the absence of exogenous expression of EGFRLs, we introduced eDHFR-cRaf into WT, dEREG, 4KO, and TKO MDCK cells generated previously (Lin et al., 2022). As expected, rapid ERK activation was observed in EREG-expressing WT and TKO (*ΔEGF, ΔHBEGF, ΔTGFA*) cells, but not in the other dEREG and 4KO cells (Fig. 4G, 4H). In short, in the confined space bounded by the tight/adherens junction, basolateral plasma membranes, and basal membrane, high-affinity EGFRLs are efficiently sequestered by the EGFR on the plasma membrane so that the low-affinity EGFRLs diffuse faster than the high-affinity EGFRLs.

### HBEGF but not EREG drives migration of confluent MDCK cells

What are the biological differences between high- and low-affinity EGFRLs expressed in MDCK cells? To investigate the effect of each EGFRL on collective cell migration, we used the previously reported “boundary assay” (Hino et al., 2020). We first seeded 4KO cells expressing EGFRL and eDHFR-cRaf into a silicone chamber and then, after removing the confinement, seeded receiver cells expressing a FRET biosensor, forming the boundary between the two cell populations (Fig. 5A). Upon the addition of m^D^cTMP, ERK waves propagated from the boundary to the receiver cells (Fig. 5B). We quantified the moving distance of each receiver cell and found that HBEGF, but not the other EGFRLs drived the receiver cells to move against the direction of ERK waves (Fig. 5C, Video S6). Previously it was report that soluble HBEGF promotes cell migration in MDCK cells, though its difference from other EGFRLs remains unclear (Singh et al., 2004). This difference could be caused by the different effects on signaling molecules other than ERK (Ronan et al., 2016). Therefore, we examined the activity changes of tyrosine kinases and ROCK using Picchu (Kurokawa et al., 2001) and Eevee-ROCK FRET biosensors (Li et al., 2017), respectively. Again, the activation waves of tyrosine kinases and ROCK were faster after EREG shedding than after HBEGF shedding (Fig. 5D, Video S7). We then reasoned that the duration and timing of ERK activation may cause a difference in the induction of cell migration. Upon EREG shedding, ERK was activated with a short delay between neighboring cells due to the fast wave propagation (Fig. 5E). In stark contrast, the peak shift of ERK activation in a manner dependent on the distance to the producer cells was obvious upon HBEGF shedding. Moreover, the full width at half maximum (FWHM) of ERK waves upon HBEGF shedding was approximately two times larger than that upon EREG shedding (Fig. 5F). Thus, the delay and/or duration of the signal may cause a difference in induced cell migration.

**Figure 5.**
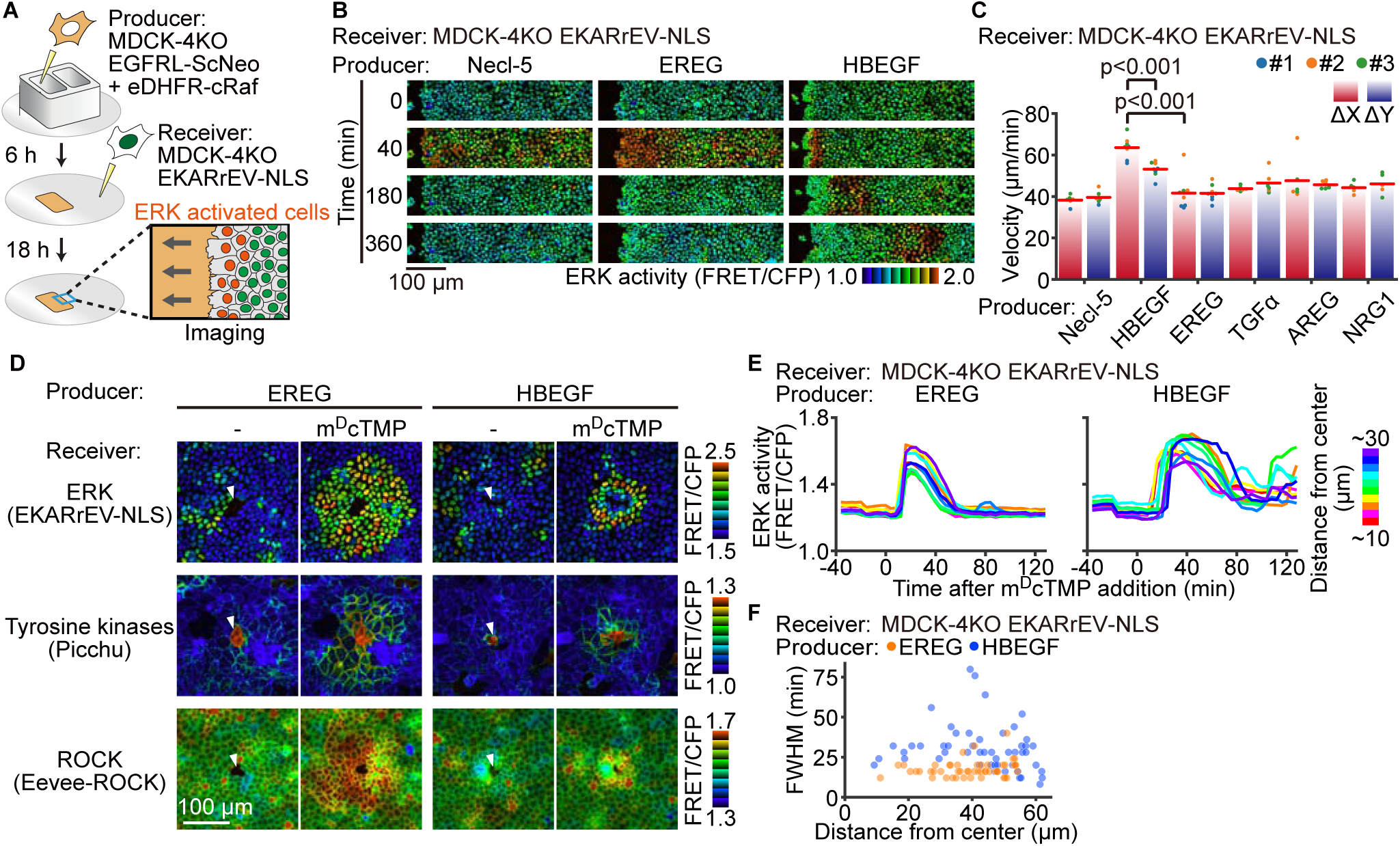
HBEGF but not EREG promotes collective cell migration. **(A)** Schematics of the boundary assay. Bath application of m^D^cTMP triggers EGFRL shedding from the producer cells, followed by the generation of ERK waves in the receiver cells. **(B)** Representative images of ERK activity (FRET/CFP) in the receiver cells adjacent to each producer cell (Video S6). m^D^cTMP was added at 0 min. **(C)** After m^D^cTMP addition, the receiver cells were tracked by the TrackMate for 10 h and the displacement was quantified. The bar graphs and red lines show the mean values. Each dot represents the average of a single field of view. n > 1000 cells from three independent experiments, depicted by the three colors. **(D)** Experiments were conducted as in Fig. 3. The receiver cells were MDCK WT cells expressing nuclear-located EKARrEV for ERK, plasma membrane-located Picchu for tyrosine kinase, or cytoplasmic Eevee-ROCK for ROCK serine/threonine kinase. White arrowheads indicate the location of EGFRL-producer cells. Images were acquired 32 min after m^D^cTMP addition (Video S7). **(E)** ERK activity (FRET/CFP) in 10 representative cells around EREG or HBEGF producers was plotted over time after m^D^cTMP addition. **(F)** Full width at half maximum (FWHM) of ERK activation in each receiver cell. n = 50 cells each from a single experiment. Statistical significance was determined by unpaired two-tailed Welch’s t-test.

### HBEGF but not EREG is sorted to lysosomes

The strength of EGFR signaling is known to be regulated by the sorting after activation (Brüggemann et al., 2021; Roepstorff et al., 2009). Thus, we tracked EGFRLs after shedding. 4KO cells expressing EREG-ScNeo or HBEGF-ScNeo were stimulated with m^D^cTMP addition and stained with anti-EEA1 and anti-Rab7 antibodies, which are early and late endosome markers, respectively (Fig. 6A). Consistent with the previous report (Roepstorff et al., 2009), we also found that EREG was localized both early and late endosomes, whereas HBEGF was localized more to the late endosomes (Fig. 6B, 6C). Since HBEGF and EREG also bind to ErbB4, we also used Erbock-ErbB1 cells, which lack all ErbB-family receptors and express human EGFR (Fig. 6D, 6E). Again, we found that HBEGF localized to late endosome more efficiently than EREG. Thus, the difference in affinity to EGFR appears to affect the fate of the EGFRL-EGFR complex within the cells and may affect the biological outcome caused by the EGFRL.

**Figure 6.**
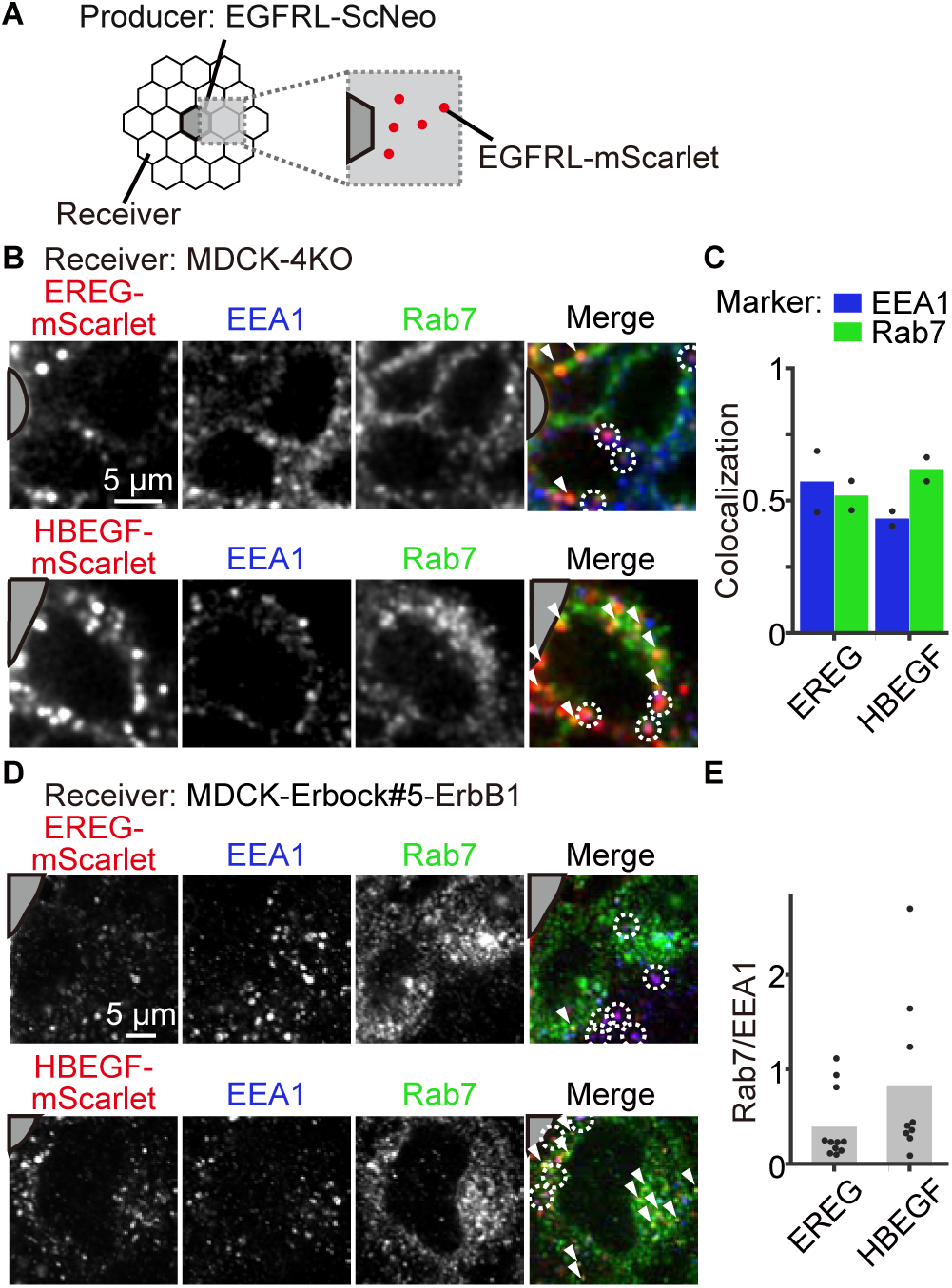
HBEGF but not EREG is sorted to late endosomes. **(A)** EGFRL-ScNeo-producer cells were co-cultured with receiver cells. After m^D^cTMP addition, cells were fixed and immunostained with endosome markers. Cells surrounding the producer cells were analyzed for colocalization of EGFRLs and the markers. **(B)** Producer cells expressing either EREG-ScNeo or HBEGF-ScNeo were co-cultured with MDCK-4KO receiver cells. Cells were fixed at 75 min after the addition of m^D^cTMP and stained with anti-EEA1 antibody and anti-Rab7 antibody. White circles and arrowheads indicate mScarlet-positive vesicles colocalized with EEA1 and Rab7, respectively. The grey area indicates the producer cells. **(C)** Fraction of mScarlet (+) vesicles colocalized with EEA1 or Rab7 from images in (B). The bar graphs show the mean values. Each dot represents the average of a single field of view. n = 2 fields of view from a single experiment. **(D)** Producer cells expressing either EREG-ScNeo or HBEGF-ScNeo were co-cultured with Erbock-ErbB1 cells, which lack all ErbB-family receptors and express human EGFR. Cells were fixed at 60 min after the addition of m^D^cTMP and stained with anti-EEA1 antibody, anti-Rab7 antibody, and anti-RFP antibody. **(E)** The proportion of mScarlet (+) vesicles colocalized with Rab7 or EEA1 from experiments in (D). The bar graphs show the mean values. Each dot represents the average of a single field of view. n = 11 (EREG), 9 (HBEGF) fields of view from three independent experiments.

### EREG is required for collective cell migration of wounded mouse epidermis

ERK waves contribute to epidermal cell migration during mouse skin wound healing (Hino et al., 2022; Hiratsuka et al., 2015). In HBEGF-deficient mice, a statistically significant delay in wound healing was reported on day 7 and day 8 (Shirakata et al., 2005). However, no statistically significant delay was reported in EREG-deficient mice (Shirasawa et al., 2004). Given the marked redundancy in the biological functions of EGFRLs, it may have been difficult to find macroscopically apparent differences in the EREG-knockout mice. Therefore, to gain insight into the function of EREG in the propagation of ERK waves, we generated EREG-knockout mouse lines expressing an ERK FRET biosensor (Komatsu et al., 2018) (Fig. S5). The mice were born with Mendelian ratios and did not show any abnormality to the detectable level, in agreement with the previous report (Shirasawa et al., 2004). We observed the auricular epidermis after wounding using two-photon microscopy as described previously (Fig. 7A) (Hiratsuka et al., 2015). Repeated waves of ERK activation were generated from the wound edge in both wild-type and *Ereg^-/-^* mice (Fig. 7B). However, the waves were extinguished at 200 µm or less in *Ereg^-/-^*mice, whereas the waves reached more than 800 µm in the WT (Fig. 7C, Video S8). Furthermore, the epidermal cells located more than 200 µm from the wound edge migrated less efficiently in *Ereg^-/-^* mice than in WT mice (Fig. 7D). Thus, we concluded that EREG serves as a long-range signal transmitter during skin wound healing.

**Figure 7.**
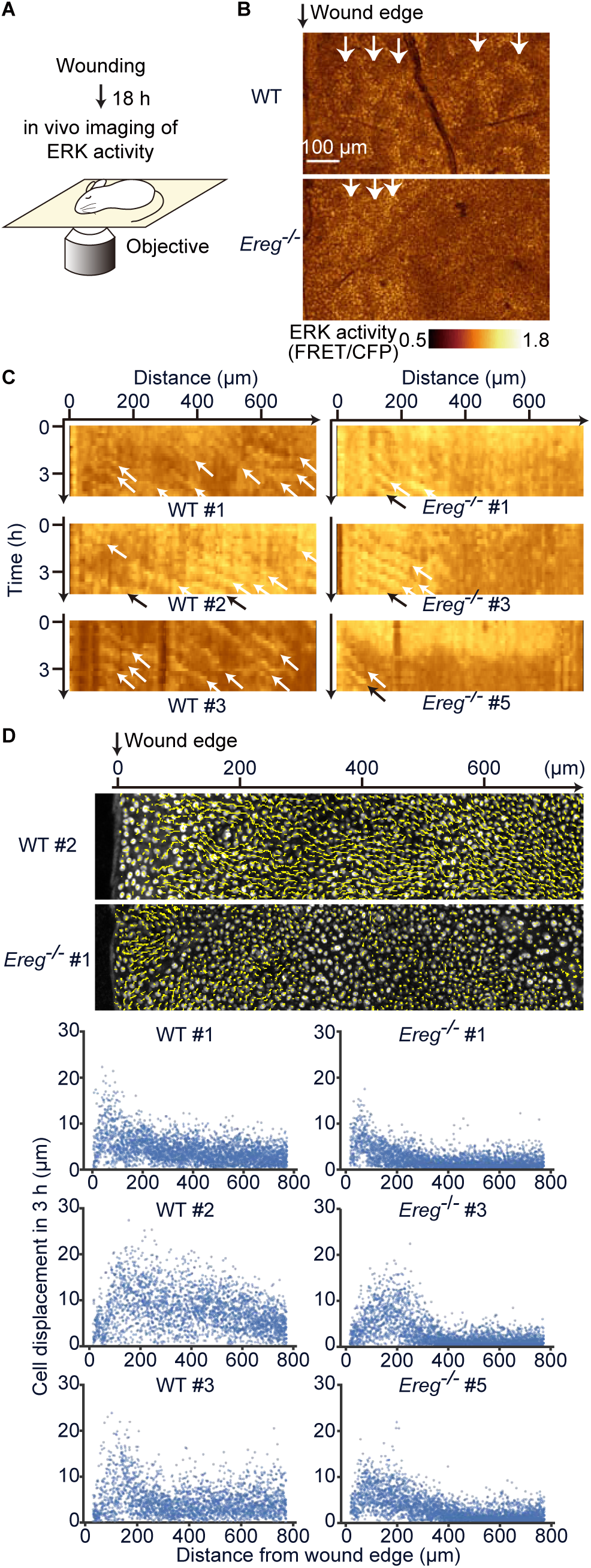
EREG is required for collective cell migration of wounded mouse epidermis. **(A)** Schematics of an *in vivo* imaging of ERK activity during wound healing of mouse ear skin expressing hyBRET-ERK-NLS. **(B)** Representative images of ERK activity (FRET/CFP) in WT or *Ereg^-/-^* mouse ear skin (Video S8). White arrows indicate ERK waves propagating from the wound edge (left black arrow). **(C)** Kymographs of ERK activity generated from time-lapse FRET/CFP ratio images. White and black arrows indicate ERK waves propagating from the wound edge (0 μm). **(D)** (Upper) Representative images of mouse skin basal layer cell trajectories for 3 h. (Bottom) Displacement of mouse skin basal layer cells toward the wound edge. Cells were tracked by the TrackMate for 3 h. Each dot represents a single cell. n > 1000 cells for each mouse.

## Discussion

Among the four EGFRLs expressed in MDCK cells, EREG, the lowest-affinity EGFRL, was found to play the major role in the propagation of ERK activation to distant cells. Intuitively we understand that the low-affinity EGFRLs reach more-distant cells than do the high-affinity EGFRLs because they will not be trapped by EGFR. However, because the low affinity dampens the efficiency of EGFR activation, we cannot foresee whether the low-affinity EGFRLs transmit signals more efficiently to distant cells than the high-affinity EGFRLs. Lauffenburger and colleagues elegantly demonstrated that decreased affinity of EGF to EGFR could increase the distance of signal propagation by theoretical and experimental approaches (DeWitt et al., 2002; Shvartsman et al., 2001); however, it has not been demonstrated in which physiological contexts the low affinity becomes an advantage for the EGFRLs. We found that the requirements are at least the intact barrier segregating the intercellular space from the apical space (Fig. 4D, 4E) as well as a physiological level of EGFR expression (Fig. 4C). Such conditions could occur at least in the skin because EREG knockout perturbed the migration of epidermal cells located distantly from the wound edge (Fig. 7). The narrow space segregated by the tight/adherens junction and basement membrane of the epidermis may render the low-affinity an advantageous property. The requirement of EREG for epithelial regeneration was also reported in the bronchiolar epithelium (Reyes et al., 2022) and intestinal mucosa (Neufert et al., 2013). In both cases, activated fibroblasts are the source of EREG; therefore, the mechanism responsible for maintaining a high concentration of EREG in the bronchiolar epithelium and intestinal mucosa may be different from that in the epidermis, but we anticipate that in all these cases the mechanism involves a restriction of the diffusion of EREG *in vivo*.

EGF is an archetype of paracrine factors, but the spatial range that EGF and other EGFRLs shed from a single cell could influence remains elusive even in tissue culture cells. The EGFRL-ScNeo probes allow us to challenge these questions. We need to consider at least three properties of EGFRLs: shedding, diffusion, and affinity to EGFR. In our present experiments we found that TGFα is most efficiently cleaved in the steady state (Fig. 1H) due to its high sensitivity to metalloproteases (Dempsey and Coffey, 1994) (Fig. 1D-1F). In agreement with this finding, Bunker et al. reported that surface expression of TGFα is almost undetectable in the absence of metalloprotease inhibitor (Bunker et al., 2021). Due to the high sensitivity to metalloproteases and high affinity to EGFR, TGFα is efficiently engulfed by essentially all cells in a culture dish (Fig. 2B).

This observation also indicates that, unlike EREG, TGFα is efficiently released to the luminal side through tight/adherens junctions. It was reported that MDCK cells cultured on permeable supports, but not on plastic dishes, shed most of TGFα to the basolateral side (Dempsey and Coffey, 1994). However, we failed to find significant differences in the distribution of TGFα between the permeable supports and cover glasses used to plate MDCK cells (Fig. S1B).

In contrast to TGFα, HBEGF was heavily accumulated inside the neighboring cells (Fig. 2B, 2H). When EGFR-deficient cells were used, HBEGF was accumulated at the basal surface of the neighboring cells (Fig. 2F), indicating that HBEGF is trapped by the heparan sulfate proteoglycans on the cell surface. Therefore, the HBEGF cleaved by metalloproteases is bound to the surface of neighboring cells through HSPGs and then taken up in an EGFR-dependent manner. In line with this view, we found that only HBEGF could activate ERK in the α-1-catenin KO cells surrounding the HBEGF-producer cells (Fig. 4D, Video S5). This property may also underpin previous observations that HBEGF functions as a juxtacrine growth factor (Higashiyama et al., 1995; Takemura et al., 1997). In this context, juxtacrine does not mean that the proHBEGF remaining uncleaved as a membrane integral protein functions to stimulate EGFR. Our observation indicates that the shed HBEGF stimulates preferentially adjacent cells simply because heparan sulfate proteoglycans decelerate HBEGF diffusion.

Why does HBEGF but not EREG induce cell movement in the boundary assay (Fig. 5A-C)? Reciprocal cycles of increase in traction force and ERK activation are the engine of collective cell migration of MDCK cells (Hirashima et al., 2023). The pulling force of the leader cells stretches the follower cells to activate EGFR and, thereby, ERK. The activated ERK not only reorganizes the actomyosin network to generate traction force but also activates ADAM17 to shed EGFRLs. EGFRLs and the pulling force cooperatively activate EGFR on the adjacent cell to ignite another cycle. To make this scenario operate, the optimal time delay between each step is critical. During collective cell migration of MDCK cells, the increase in traction force is followed by ERK activation with a 2 min delay (Hino et al., 2020). ERK activation is followed by accumulation of phosphorylated myosin light chain with a 6 min delay(Aoki et al., 2017). It takes 7 min for an ERK wave to pass through a single cell (Hirashima et al., 2023). This delay enables cells to move in a peristaltic manner. In fact, an analysis using optogenetic tools found that ERK activity waves at a velocity of 2.0–3.0 μm/min are optimal for driving MDCK cells (Aoki et al., 2017). The velocity of the ERK activity wave generated by HBEGF, 1.0 µm/min, is slower than the optimal value (Fig. 3H), but this may be within a permissible range because the ERK wave velocity in mouse epidermis is 1.4 μm/min (Hiratsuka et al., 2015). Meanwhile, EREG-generated waves propagating at 4 µm/min may be moving too fast to drive MDCK cell movements.

Other factors that might have caused the phenotypical difference between HBEGF and EREG are the duration of ERK activation and the different sorting pathways after binding to EGFR. With respect to the first factor, the full width at half maximum (FWHF) of ERK waves upon HBEGF shedding was approximately two-fold larger than that upon EREG shedding (Fig. 5F). In support of this finding, in the case of migration of MCF7 driven by bath application of EGF, low dose, but not high dose EGF can induce cell migration because of sustained ERK activation (Brüggemann et al., 2021). It has been reported that different ligands undergo different endocytic sorting(Roepstorff et al., 2009). In line with this, we found that HBEGF but not EREG was sorted into late endosomes (Fig. 6). Thus, the intracellular signaling pathways may differ between HBEGF and EREG. This issue should be examined in future studies.

Chemogenetic tools are widely used to perturb intracellular signaling cascades, but much less frequently used to untangle intercellular communications. Here we employed the SLIPT system to activate ADAM17 (Suzuki et al., 2022), thereby shedding EGFRLs. This approach will also be useful for studying the effects of other ADAM17 target molecules such as TNFα. On the other hand, this method is not specific in that all targets of ERK will be activated in cells with the SLIPT system, and all ADAM17 target molecules will be cleaved. The cleavage of the ADAM17 target is not necessarily dependent only on ADAM17. For example, cleavage of EGFRLs is also regulated by iRhoms (Li et al., 2015; Tang et al., 2020). ERK is known to activate ADAM17 (Fan and Derynck, 1999), but other kinases can also modulate ADAM-family metalloproteases (Dang et al., 2013; Miller et al., 2013), raising a question about the specificity. Moreover, we failed to activate ADAM10 in MDCK cells, which prevented us from studying the roles of EGF and BTC. This should be overcome in the future.

In conclusion, we have revealed that low-affinity EGFRLs propagate ERK activation faster and further *in vitro* and *in vivo*. Our findings will shed light on the importance of low-affinity ligands in cell-to-cell communication in the physiological context, thus bringing us closer to understanding the significance of the existence of multiple EGFRLs.

## Limitations of this research

The effect of fusion proteins has not been thoroughly tested partly due to the availability of antibodies against EGFRLs. The subcellular distribution, sensitivity to ADAM17, and biological effect as the EGFR agonist, of the EGFRLs-ScNeos generally agree with previous reports, but further examination will be required. We failed to induce the cleavage of EGF-ScNeo and BTC-ScNeo, which are expected to be substrates of ADAM10. We applied calcium ionophore or other stimuli that are reported to activate ADAM10, but we did not observe any changes in the fluorescence ratio.

## Acknowledgments

We are grateful to the members of the Matsuda Laboratory for their helpful input, to K. Hirano, T. Uesugi, and K. Takakura, who provided technical assistance, and to the Medical Research Support Center of Kyoto University. This work was supported by the Kyoto University Live Imaging Center. Financial support was provided by JSPS KAKENHI grants (21H05226 to K.T., and 19H00993 and 20H05898 to M.M.), a JST Moonshot R&D grant (JPMJPS2022 to M.M.), NIH grants (GM148363 and CA089151 to A.S.) and a JST SPRING grant (JPMJSP2110 to E. D.).

## AUTHOR INFORMATION

## Author contributions

Conceptualization, Methodology, Validation, Formal Analysis: E.D., M.M., K.T.; Investigation: E.D., S.L., K.M.; Data Curation: E.D., M.M., and K.T.; Resources: A.T., K.S., S.T., T.O., M.F., and A.D.S.; Writing – Original Draft: E.D.; Writing – Review & Editing: E.D., A.S., M.M., K.T.; Supervision: M.M. and K.T.; Project Administration: M.M. and K.T.; Funding Acquisition: E.D., M.M., and K.T.

## Methods

### Plasmids

The following cDNAs encoding human EGFRLs were used to construct EGFRL-ScNeo plasmids: pro-EGF (Biological Resource Center, National Institute of Technology and Evaluation, Tokyo, Japan; account no. AK299306), pro-HBEGF and pro-TGFα (gifts from Ryo Iwamoto, Osaka University, Osaka, Japan), pro-EREG (NCBI CCDS database no. CCDS3564.1), pro-AREG (NCBI CCDS database no. CCDS3565.1), pro-EPGN (NCBI CCDS database no. CCDS59478.1), pro-NRG1 (NCBI CCDS database no. CCDS6083.1), and pro-BTC (NCBI CCDS database no. CCDS3566.1). The sequences obtained from the NCBI CCDS database were used to synthesize cDNAs by GeneArt (Thermo Fisher Scientific, Waltham, MA). The cDNA of mScarlet was obtained from Addgene (Addgene no. 85044). The cDNA of mNeonGreen(Shaner et al., 2013) was synthesized with codon optimization by GeneArt (Thermo Fisher Scientific). Expression plasmids of EGFRL-ScNeo were constructed according to the standard PCR-based amplification. An HA (YPYDVPDYA) tag, a flexible linker (GGGSGGGS), mScarlet, and mNeonGreen were inserted into the cDNA sequences of human EGFRLs (Fig. 1B). The expression plasmids and established cell lines are summarized in Supplementary Tables S1A and S1B. Briefly, the mScarlet cDNA was inserted at the 5’ end of the EGF domain of HBEGF, AREG, BTC and EPGN, so that the protease cleave site of propeptide was at the 5’ end of mScarlet cDNA. cDNA of HA-tag, mScarlet and the flexible linker were fused together in this order and inserted at the 5’ end of the EGF domain of EGF, TGFα and EREG. The mNeonGreen cDNA was inserted into the 3’ end of the EGFRL cDNAs except in the case of TGFα, for which the last 6 nucleotides were modified to encode valines. The cDNAs encoding the fusion proteins, collectively called EGFRL-ScNeo, were subcloned into the pCSII vector (Miyoshi et al., 1998) and pPB vector (Yusa et al., 2009) using Ligation high Ver. 2 (TOYOBO, Osaka, Japan) or the In-Fusion cloning kit (Clontech, Mountain View, CA). To generate pCSIIpuro-Necl5-ScNeo, cDNAs coding the signal peptide of Necl5 (a.a. 1–28), mScarlet, Necl5 (a.a. 29–408), and mNeonGreen were obtained from the previously established plasmid pPBbsr2-Necl5-ScNeo (Ichise et al., 2022) and assembled into pCSII vectors by using Ligation high Ver. 2 (TOYOBO). pPBpuro-EGFP-eDHFR(69K6)-cRaf (Addgene no. 178849) was reported previously (Suzuki et al., 2022). cDNA of miRFP703 was a gift from Kazuhiro Aoki (Kyoto University, Kyoto, Japan). Expression plasmids encoding human EGFR were constructed previously (Matsuda et al., 2023). Expression plasmids encoding FRET ERK biosensors were reported previously (Kawabata and Matsuda, 2016; Lin et al., 2022). An expression plasmid encoding FRET ROCK biosensors was reported previously (Hino et al., 2020). An expression plasmid encoding the FRET tyrosine kinase biosensor was subjected to W169L mutation from the original version (Hino et al., 2022). An expression plasmid encoding the FRET biosensor for ADAM17, pPBbsr2-TSen, was developed according to the previous report (Chapnick et al., 2015). The plasmids used in this paper are listed in STAR Methods.

### Reagents and antibodies

The following reagents were used: dimethyl sulfoxide (no. 13445-74; Nacalai Tesque, Kyoto, Japan), 12-O-tetradecanoylphorbol 13-acetate (TPA) (no. P-1680; LC Laboratories, Woburn, MA), Marimastat (no. SC-202223; Santa Cruz Biotechnology, Dallas, TX), EGF (no. E9644; Sigma-Aldrich, St. Louis, MO), HBEGF (no. 100-47; PeproTech, Cranbury, NJ), EREG (no. 100-04; PeproTech), TGFα (no. 239-A-100; R&D Systems, Minneapolis, MN), trametinib (no. T-8123; LC Laboratories), Cytochalasin D (no. 250255; Calbiochem, La Jolla, CA), and bovine serum albumin (no. A2153; Sigma-Aldrich). m^D^cTMP was synthesized as described previously (Nakamura et al., 2020).

The following primary and secondary antibodies were used for immunoblotting: anti-mCherry rabbit antibody (no. ab167453; Abcam, Cambridge, UK; 1:1,000 dilution); anti-mNeonGreen rabbit antibody (no. 53061; Cell Signaling Technology, Beverly, MA; 1:1,000 dilution); anti-phospho-EGFR (Tyr1068) rabbit antibody (no. 3777; Cell Signaling Technology; 1:1,000 dilution); anti-alpha Tubulin mouse antibody (DM1A) (no. 62204; Thermo Fisher Scientific; 1:1,000 dilution); IRDye 680-conjugated goat anti-mouse IgG antibody (no. 926-32220; LI-COR Biosciences, Lincoln, NE; 1:10,000 dilution); and IRDye 800CW goat anti-rabbit IgG antibody (no. 926-32211; LI-COR Biosciences; 1:10,000 dilution).

The following primary and secondary antibodies were used for immunofluorescence: anti-GP135 mouse antibody (no. MABS1327; Merck Millipore, Burlington, MA; 1:100 dilution); anti-ZO-1 rabbit antibody (no. 40-2200, Thermo Fisher Scientific, Waltham, MA; 1:100 dilution); anti-EEA1 mouse antibody (no. 610457; BD Biosciences; 1:250 dilution); anti-Rab7 rabbit antibody (no. 9367; Cell Signaling Technology; 1:100 dilution); anti-RFP rat antibody (no. 5f8; chromotek; 1:500 dilution); AMCA-conjugated donkey anti-mouse IgG (H+L) antibody (no. 715-155-151; Jackson ImmunoResearch, West Grove, PA; 1:25 dilution); Cy5-conjugated donkey anti-rabbit IgG (H+L) antibody (no. 711-175-152; Jackson ImmunoResearch; 1:250 dilution); Alexa 405-conjugated goat anti-mouse IgG (H+L) antibody (no. A-31553; Thermo Fisher Scientific; 1:250 dilution); Alexa 647-conjugated goat anti-rabbit IgG (H+L) antibody (no. A-21245; Thermo Fisher Scientific; 1:100 dilution); Alexa 546-conjugated goat anti-rat IgG (H+L) antibody (no. A-11081; Thermo Fisher Scientific; 1:500 dilution).

### Cell culture

MDCK and Lenti-X 293T cells were provided by RIKEN BioResource Center (no. RCB0995) and Clontech (no. 632180), respectively. MDCK II and Claudin quinKO cells were reported previously (Otani et al., 2019). These cells were cultured in DMEM (no. 044-29765; Wako, Osaka, Japan) supplemented with 10% FBS (F7524; Sigma-Aldrich), 100 units mL^-1^ penicillin, and 100 μg mL^-1^ streptomycin (no. 26253-84; Nacalai Tesque) in a 5% CO2 humidified incubator at 37°C.

### Establishment of stable cell lines

A lentiviral expression system was employed to establish MDCK cells stably expressing EGFRL-ScNeo as described previously (Hino et al., 2020; Lin et al., 2022). Briefly, for the preparation of the lentivirus, pCSII-EGFRL-ScNeo, a vector for lentiviral transduction (Miyoshi et al., 1998), psPAX2 (Addgene Plasmid: no. 12260), and pCMV-VSV-G-RSV-Rev were co-transfected into Lenti-X 293T cells by using polyethyleneimine (no. 24765-1; Polyscience Inc., Warrington, PA). MDCK cells were incubated with the lentivirus and after 2 days of incubation, the cells were treated with 2 μg mL^-1^ puromycin (no. P-8833; Sigma-Aldrich), 10 μg mL^-1^ blasticidin S (no. 029-18701; Wako) or 100 µg mL^-1^ zeocin (no. 11006-33-0; InvivoGen, San Diego, CA) for the selection. The obtained MDCK cells were sorted using an FACS Aria IIu cell sorter (Becton Dickinson, Franklin Lakes, NJ) with mNeonGreen fluorescence to achieve a uniform expression level of the EGFRL-ScNeo. MDCK cells stably expressing EGFRL-ScNeo, miRFP703-eDHFR(69K6)-cRaf, TSen, human EGFR or tyrosine kinase biosensor were established with a piggyBac transposon system. pPB plasmids and pCMV-mPBase(neo-) encoding piggyBac transposase (Yusa et al., 2009) were co-transfected into MDCK cells by electroporation with an Amaxa nucleofector (Lonza, Basel, Switzerland), followed by selection with 2 μg mL^-1^ puromycin (no. P-8833; Sigma-Aldrich), 10 μg mL^-1^ blasticidin S (no. 029-18701; Wako) or 100 µg mL^-1^ zeocin (no. 11006-33-0; InvivoGen). MDCK cells stably expressing EKARrEV-NLS (Lin et al., 2022), EKAREV-NLS (Kawabata and Matsuda, 2016), and Eevee-ROCK-NES (Hino et al., 2020) were reported previously. The cell lines used in this paper are listed in Table S1B.

### CRISPR/Cas9-mediated KO cell lines

For CRISPR/Cas9-mediated KO of genes encoding EGFRLs, ErbB receptors, ADAM17, α-1-catenin, E-cadherin, and p120-catenin, single guide RNAs (sgRNA) targeting the exons were designed using CRISPRdirect (Naito et al., 2015) as described previously (Hino et al., 2022; Lin et al., 2022; Matsuda et al., 2023). The gRNA sequences are listed in Table S1C. Oligonucleotide DNAs for the sgRNA were cloned into lentiCRISPRv2 (Addgene no. 52961) (Sanjana et al., 2014), pX458 (Addgene no. 48138) (Ran et al., 2013) or pX459 (Addgene no. 62988) (Ran et al., 2013). The expression plasmids for sgRNA and Cas9 were introduced into MDCK cells by lentiviral infection or electroporation. For lentivirus production, the lentiCRISPRv2-derived expression plasmid, psPAX2 (Addgene no. 12260) and pCMV-VSV-G-RSV-Rev (Miyoshi et al., 1998) were co-transfected into Lenti-X 293T cells using polyethylenimine (no. 24765-1; Polyscience Inc.). The infected cells were selected with media containing the following antibiotics, depending on the drug resistance genes carried by the lentiCRISPRv2-derived plasmids: 100 µg mL^-1^ zeocin (no. 11006-33-0; InvivoGen), 2.0 µg mL^-1^ puromycin (no. P-8833; Sigma-Aldrich), 200 μg mL^-1^ hygromycin (no. 31282-04-9; Wako) and/or 800 μg mL^-1^ neomycin (no. 16512-52; Nacalai Tesque). For electroporation, pX458 or 459-derived expression plasmids were transfected into MDCK cells by an Amaxa Nucleofector II (Lonza). The transfected cells were selected with 2.0 µg mL^-1^ puromycin. After the selection, reduction in expression levels of the E-cadherin and p120-catenin proteins was confirmed by immunoblotting. Bulk cells were used for the experiments. After the selection and subsequent single-cell cloning, reduction in expression levels of α-1-catenin protein was confirmed by immunoblotting. For the KO of EGFRLs, ErbB receptors, and ADAM17, after single-cell cloning, genomic DNAs were isolated with SimplePrep reagent (no. 9180; TAKARA bio, Kusatsu, Japan) according to the manufacturer’s instructions. PCR was performed using KOD FX neo (no. KFX-201; TOYOBO) for amplification with the designed primers, followed by DNA sequencing.

### Fluorescence imaging with a confocal laser microscope

Cells were observed with a Leica TCS SP8 FALCON confocal microscope (Leica-Microsystems, Wetzlar, Germany) equipped with an HC PL APO 40x/1.30 OIL CS2 objective, an HC PL APO 63x/1.40 OIL CS2 objective, Lecia HyD SMD detectors, a white light laser of 80 MHz pulse frequency, a Diode 405 (VLK 0550 T01; LASOS, Jena, Germany), a 440 nm diode laser (PDL 800-D; PicoQuant, Berlin, Germany), and a stage top incubator (Tokai Hit, Fujinomiya, Japan) to maintain 37°C and 5% CO2. The following excitation wavelengths and emission band paths were used for the imaging: for CFP and YFP imaging, 440 nm excitation, 467–499 nm and 520–550 nm emission, respectively; for mNeonGreen imaging, 505 nm excitation, 515–560 nm emission; for mScarlet imaging, 569 nm excitation, 579–650 nm emission; for miRFP imaging, 670 nm excitation, 680–800 nm emission; for Alexa 405 imaging, 405 nm excitation, 415– 500 nm emission; for Alexa 546 imaging, 561 nm excitation, 570-640 nm emission; for Alexa 647 imaging, 650 nm excitation, 660–750 emission. To eliminate the background signal, the time gate for mScarlet and miRFP fluorescence detection was set from 0.5 ns to 6.5 ns and 0.3 ns to 6.0 ns, respectively.

### Time-lapse imaging with wide-field fluorescence microscopes

Wide-field fluorescence images were acquired essentially as described previously (Aoki and Matsuda, 2009). Briefly, cells cultured on glass-bottom plates were observed under an ECLIPSE Ti2 inverted microscope (Nikon, Tokyo, Japan) equipped with a 10X/0.30 PlanFluor, a 20X/0.70 S Plan Fluor LWD ADM, a 40X/0.60 S Plan Fluor ELWD ADM, an ORCA Fusion Digital CMOS camera (Hamamatsu Photonics K.K., Hamamatsu, Japan), an X-Cite TURBO LED light source (Excelitas Technologies, Waltham, MA), a Perfect Focus System (Nikon), a TI2-S-SE-E motorized stage (Nikon), and a stage top incubator (Tokai Hit) to maintain 37°C and 5% CO2. The following filters were used for the time-lapse imaging: for CFP and YFP imaging, a 434/32 excitation filter (Nikon), a dichroic mirror 455 (Nikon), and 480/40 and 535-30 emission filters (Nikon) for CFP and YFP, respectively; for mScarlet imaging, a 570/40 (Nikon) excitation filter, a dichroic mirror 600 (Nikon), and a 645/75 emission filter (Nikon); for miRFP703 imaging, an FF01-640/14 excitation filter (Semrock, Rochester, NY), a dichroic mirror 660 (Nikon), and a 700/75 emission filter (Nikon).

### Image processing for the FRET/CFP ratio

Image processing for FRET/CFP ratio images was performed with Fiji (Schindelin et al., 2012). The background intensity was subtracted by using the subtract-background function and subsequently processed with a median filter to reduce noise. The processed images were subjected to image calculation and the ratio values were binned into 8 steps to obtain 8-color FRET/CFP ratio images. To convey the brightness of the original images to the FRET/CFP ratio images, the 8-color FRET/CFP ratio images were multiplied by the corresponding intensity-normalized grayscale image.

### Fluorescence imaging and mScarlet/mNeonGreen ratio quantification of EGFRL-ScNeo

For fluorescence imaging of EGFRL-ScNeo, 2.0 x 10^4^ MDCK-EGFRL-ScNeo cells were seeded on a 96-well glass-bottom plate (AGC TECHNO GLASS, Shizuoka, Japan) coated with 0.3 mg mL^-1^ type I collagen (Nitta Gelatin, Osaka, Japan). The medium was replaced with DMEM supplemented with 10% FBS, 100 units mL^-1^ penicillin, and 100 μg mL^-1^ streptomycinstreptomycin on Day 1 and observed on Day 2 under a TCS SP8 microscope at a resolution of 0.045 μm/pixel using a 63x/1.40 NA objective, 505 nm excitation for 1.5% and 569 nm for 2.0% of a white light laser, and HyD SMD detectors for 515–560 nm and 579–650 nm for gain of 60%, respectively.

For quantification of the mScarlet/mNeonGreen ratio at the plasma membrane, the following commands were applied sequentially using Fiji: “8-bit,” “Make Binary,” “Watershed,” “Open,” “Close-,” and “Analyze Particles…,” “size=1-Infinity circularity=0.00–0.60” for making ROIs on the cell membrane. Intensities of mScarlet and mNeonGreen were measured for the mScarlet/mNeonGreen ratio.

### Western blotting analysis of EGFRL-ScNeo

For western blotting of cell lysates, 5.0 x 10^5^ MDCK-EGFRL-ScNeo cells were plated in a 6-well plate (no. 140675; Thermo Fisher Scientific). One day after seeding, cells were lysed with SDS sample buffer containing 62.5 mM Tris-HCl (pH 6.8), 12% glycerol, 2% SDS, 40 ng mL^-1^ bromophenol blue, and 5% 2-mercaptoethanol, followed by sonication with a Bioruptor UCD-200 (Cosmo Bio, Tokyo, Japan). After boiling at 95°C for 5 min, the samples were separated by SDS-PAGE on SuperSep Ace 5–20% precast gels (Wako), and transferred to polyvinylidene difluoride membranes (Merck Millipore, Burlington, MA) for Western blotting. All antibodies were diluted in Odyssey blocking buffer (LI-COR Biosciences). Proteins were detected by an Odyssey Infrared Imaging System (LI-COR Biosciences). Cleaved/total pro-EGFRL was calculated by dividing the sum of the intensities of mNeonGreen bands corresponding to the cleaved form by the sum of the total mNeonGreen bands.

### Quantification of EGFRL in the culture supernatant

The number of EGFRL molecules liberated into the medium was calculated by Western blotting analysis of mScarlet. For the collection of the culture supernatant, 2.5 x 10^6^ MDCK-4KO-EGFRL-ScNeo cells were plated in a 10 cm dish (no. 150466; Thermo Fisher Scientific). One day after seeding, cells were washed with PBS and the medium was replaced with DMEM supplemented with 100 units mL^-1^ penicillin and 100 μg mL^-1^ streptomycin. Twenty-four hours later, the medium was collected and centrifuged at 200 g for 3 min to remove cells. Supernatants were mixed with a 1/5th volume of 6xSDS sample buffer. After boiling at 95°C for 5 min, the samples were separated by SDS-PAGE as described above. The amount of mScarlet in each sample was quantified from Western blotting with anti-mCherry antibody and the calibration by mScarlet.

For the calibration of mScarlet, ArcticExpress (DE3) Competent Cells (no. 230192; Agilent Technologies, Santa Clara, CA) were transformed with an IPTG-inducible mScarlet expression plasmid, pRSETB-mScarlet. Cells were cultured in LB medium supplemented with IPTG 1 mM by shaking. The cells were then collected and centrifuged at 15,000 x g for 1 min. The precipitate was mixed with 1xSDS sample buffer. After boiling at 95°C for 5 min, the samples were separated by SDS-PAGE as described above. The gel was stained with CBB, and the amount of mScarlet in the sample was calculated using a standard curve of BSA.

### Analysis of ADAM sensitivity

For the time-lapse imaging of EREG-ScNeo, 4.0 x 10^4^ MDCK-EREG-ScNeo cells were seeded on a 96-well glass-bottom plate (Matsunami Glass Ind. Ltd., Kishiwada, Japan) coated with 0.3 mg mL^-1^ type I collagen. The medium was DMEM supplemented with 10% FBSFBS, 100 unit mL^-1^ penicillin, and 100 μg mL^-1^ streptomycin. One day after seeding, cells were observed under a TCS SP8 microscope at a resolution of 0.18 μm/pixel using a 63x/1.40 NA objective, 505 nm excitation for 2.5% and 569 nm for 4.0% of a white light laser, and HyD SMD detectors for 515–560 nm and 579–650 nm for gain of 60%, respectively. During observation, TPA was added to 10 nM and Marimastat was added to 10 μM. mScarlet/mNeonGreen ratio images were generated using the Fiji plug-in. Pseudo-color ratio images were generated by multiplying 8-color mScarlet/mNeonGreen images with the corresponding grayscale images.

For the quantification of the mScarlet/mNeonGreen ratio of EGFRL-ScNeo, 2.0 x 10^4^ MDCK-EGFRL-ScNeo cells were seeded on a 96-well glass-bottom plate (Matsunami Glass). The medium was DMEM supplemented with 10% FBSFBS, 100 unit mL^-1^ penicillin, and 100 μg mL^-1^ streptomycin. For ADAM activation, one day after seeding, cells were observed under a TCS SP8 microscope at a resolution of 0.18 μm/pixel using a 63x/1.40 NA objective, 505 nm excitation for 4.0% and 569 nm for 6.0% of a white light laser, and HyD SMD detectors for 515–560 nm and 579–650 nm for gain of 60%, respectively. During observation, TPA was added to 10 nM. For ADAM inhibition, 3 h after seeding, cells were supplemented with Marimastat 10 μM or DMSO 0.1%. One day after seeding, cells were observed under a TCS SP8 microscope at a resolution of 0.28 μm/pixel using a 40x/1.30 NA objective, 505 nm excitation for 0.7% and 569 nm for 1.0% of a white light laser, and HyD SMD detectors for 515–560 nm and 579–650 nm for gain of 60%, respectively. The mScarlet/mNeonGreen ratio at the plasma membrane was quantified as described above.

### Analysis of ERK activation by the supernatant of EGFRL-ScNeo

For the collection of supernatants of EGFRL-ScNeo-expressing cells, 2.0 x 10^4^ MDCK-4KO-EGFRL-ScNeo cells were seeded on a 96-well plate. Five hours after seeding, cells were washed with PBS and the medium was replaced with Medium 199 (11043023; Life Technologies, Carlsbad, CA) supplemented with 100 units mL^-1^ penicillin and 100 µg mL^-1^ streptomycin. Twenty hours after the medium change, the supernatant was collected. The number of ligand molecules in the supernatant was normalized with the amount of mScarlet by Western blotting as described above.

For the detection of ERK activity by live imaging, 1.0 x 10^4^ MDCK-4KO-EKARrEV-NLS cells were seeded on a 96-well glass-bottom plate (Matsunami Glass) coated with 0.3 mg mL^-1^ type I collagen. One day after seeding, the medium was replaced with Medium 199 supplemented with 100 units mL^-1^ penicillin and 100 µg mL^-1^ streptomycin. After 2 h, cells were observed under an ECLIPSE Ti2 microscope.

During observation, the medium was replaced with supernatants of MDCK-4KO-EGFRL-ScNeo or MDCK-4KO-EKARrEV-NLS. Image processing for FRET/CFP ratio images was performed with Fiji essentially as described previously (Lin et al., 2022). Briefly, cells were tracked using the Fiji TrackMate plugin (Tinevez et al., 2017) to measure the time course of the FRET/CFP ratio in each cell. The time-series data of the coordinates of each cell and the FRET/CFP ratio representing ERK activity were processed by using MATLAB. The FRET/CFP ratio in each cell was normalized with the average FRET/CFP ratios of 12 timepoints before the replacement of supernatants.

For the detection of ERK activity by western blotting, 2.0 x 10^4^ MDCK-4KO-EKARrEV-NLS cells were seeded on a 96-well glass-bottom plate (Matsunami Glass) coated with 0.3 mg mL^-1^ type I collagen. One day after seeding, the medium was replaced with Medium 199 supplemented with 100 units mL^-1^ penicillin and 100 µg mL^-1^ streptomycin. After 48 h, the medium was replaced with supernatants of MDCK-4KO-EGFRL-ScNeo or MDCK-4KO-EKARrEV-NLS. 10 minutes after the medium change, cells were lysed with SDS buffer, followed by SDS-PAGE as described above. For the controls of the experiment, MDCK-4KO-EKARrEV-NLS cells were supplemented with EGF 10 ng mL^-1^ for 10 min or Trametinib 200 nM for 30 min.

### Co-culture experiment of EGFRL-ScNeo

For the co-culture experiment of EGFRL-ScNeo, 1.0 x 10^2^ MDCK-EGFRL-ScNeo cells were mixed with 4.0 x 10^4^ parental MDCK, MDCK-Erbock#5, or MDCK-dErbB1#1 cells and seeded on a 96-well glass-bottom plate (Matsunami Glass) coated with 0.3 mg mL^-1^ type I collagen. One day after seeding, the medium was replaced with Medium 199 supplemented with 100 unit mL^-1^ penicillin and 100 μg mL^-1^ streptomycin. The cells were observed under a TCS SP8 microscope at a resolution of 0.57 μm/pixel using a 40x/1.30 NA objective, 505 nm excitation for 1.5% and 569 nm for 8.0% of a white light laser, and HyD SMD detectors for 515–560 nm for gain of 80% and 579–650 nm for gain of 150%. Z stack images were acquired every 1 μm for 21 slices.

For quantification of mScarlet signals around producer cells, the following commands of Fiji were applied sequentially on mScarlet and mNeonGreen z stack images: “Median…,” “radius=3 stack,” “Z Project…,” “projection=[Average Intensity]”. For making ROIs on producer cells, the mNeonGreen images were further processed as follows: “Make Binary,” with Li method, “Open,” “Dilate”. For quantification of surrounding mScarlet signals, the ROI was dilated every 5 pixels, and the difference in mScarlet intensities between ROIs was measured. mScarlet intensities were normalized with mNeonGreen intensities of producer cells. Then the square root of the normalized mScarlet intensities was plotted against the distance from the first ROI.

The 3D image reconstruction was performed by using Volocity software (PerkinElmer, Waltham, MA).

### Flow cytometry analysis of co-culture experiments

MDCK-EGFRL-ScNeo cells and parental MDCK cells were mixed at a cell number ratio of 1:1 to 1:400 in total 2.0 x 10^5^ cells and seeded on a 12-well plate (no. 150628; Thermo Fisher Scientific). One day after seeding, cells suspended in PBS containing 3% FBS were analyzed with a FACS Aria IIu cell sorter (Becton Dickinson). The following combinations of lasers and emission filters were used for the detection of fluorescence: for mNeonGreen, a 488-nm laser, and a DF530/30 filter; for mScarlet, a 561-nm laser and a DF610/20 filter (Omega Optical, Brattleboro, VT). Cells were gated for size and granularity to exclude cell debris and aggregates. Data analysis was performed using FlowJo software (Tree Star, Ashland, OR).

### Fluorescence imaging of TSen

For the imaging of MDCK-TSen cells expressing miRFP703-eDHFR(69K6)-cRaf, 4.0 x 10^4^ cells were seeded on a 96-well glass-bottom plate (Matsunami Glass) coated with 0.3 mg mL^-1^ type I collagen. One day after seeding, the medium was replaced with Medium 199 supplemented with 100 units mL^-1^ penicillin and 100 µg mL^-1^ streptomycin. After 2 h, cells were observed under a TCS SP8 microscope at a resolution of 0.18 μm/pixel using a 63x/1.40 NA objective, 440 nm excitation of a diode laser, and HyD SMD detectors for 467–499 nm and 520–550 nm for gain of 80%, respectively. During observation, m^D^cTMP was added to 0.01 to 10 μM or DMSO 0.1%.

For quantification of the FRET/CFP ratio at the plasma membrane, the following commands were applied sequentially: “8-bit,” “Make Binary,” “Watershed,” “Open,” “Close-,” and “Analyze Particles…,” “size=1-Infinity circularity=0.00–9.60” for making ROIs on the cell membrane. Intensities of CFP and FRET were measured for the FRET/CFP ratio.

### Shedding of EGFRL and observation of ERK activity by SLIPT

MDCK-4KO-EGFRL-ScNeo cells or MDCK-EKARrEV-NLS cells with or without EGFRL gene knockout expressing miRFP703-eDHFR(69K6)-cRaf were used as the producer cells. MDCK-EKARrEV-NLS cells with or without gene knockout of ErbB receptors or adherens junction molecules, or MDCK II-EKARrEV-NLS cells, Claudin quinKO-EKARrEV-NLS cells, MDCK-5102HRasCT(Picchu) cells, and MDCK-Eevee-ROCK-NES cells were used as the receiver cells. For the imaging of the SLIPT assay, 1.0 x 10^2^ producer cells were mixed with 8.0 x 10^4^ receiver cells and seeded on a 96-well glass-bottom plate (Matsunami Glass) coated with 0.3 mg mL^-1^ type I collagen. Three hours after seeding, the medium was replaced with DMEM supplemented with 1% BSA, 100 unit mL^-1^ penicillin, and 100 μg mL^-1^ streptomycin. One day after seeding, the medium was replaced with Medium 199 supplemented with 100 unit mL^-1^ penicillin and 100 μg mL^-1^ streptomycin. After 2 h, cells were imaged under an ECLIPSE Ti2 microscope or a TCS SP8 microscope. For the ECLIPSE Ti2 microscope, a 20X/0.70 NA or 40X/0.60 NA objective was used at a resolution of 1.3 μm/pixel or 0.65 μm/pixel, respectively. For the TCS SP8 microscope, a 40x/1.30 OIL CS2 objective was used at a resolution 0.76 μm/pixel, 440 nm excitation of a diode laser, and HyD SMD detectors for 467–499 nm for gain of 80% and 520–550 nm for gain of 40%, respectively. During observation, m^D^cTMP was added to 10 μM.

### Quantification of the velocity of the ERK activity wave

The velocity of the radial ERK activity wave was analyzed using MATLAB (MathWorks, Natick, MA) as described previously with a slight modification (Hiratsuka et al., 2015). First, on the CFP image, nuclei were automatically recognized with a segmentation program. With each nucleus, the FRET/CFP ratio was calculated for all time frames. Second, the FRET/CFP values of each nucleus were smoothed by 20-min moving averages. After noise reduction with the Savitzky–Golay filter, the peak time, i.e., the time when the FRET/CFP ratio reaches the zenith, was obtained. Third, for each nucleus of the receiver cells, the distance from the center was plotted against the peak time. Then the velocity of ERK propagation was approximated by a linear model.

### Quantification of the radius of the ERK activity wave

The radius of the ERK activity wave was analyzed using Python as described previously with a slight modification (Watabe et al., 2023). Ratio images of MDCK cells expressing EKARrEV-NLS were created after background subtraction. A median filter and a Gaussian 2D filter were applied to each image for noise reduction. The ratio image was normalized by a minimum intensity projection along the time axis. The processed images were binarized with a predetermined threshold and processed by morphological opening and closing to refine the ERK-activated area. Center coordinates and equivalent circle radii were obtained from each ERK-activated area. The maximum radius of the equivalent circle was defined as the radius of the ERK activity wave.

### Boundary assay

Cells were seeded and observed under microscopy as described previously (Hino et al., 2020). Briefly, MDCK-4KO-EGFRL-ScNeo cells expressing miRFP703-eDHFR(69K6)-cRaf were seeded in a well of a Culture-Insert 2 well (no. 81176; ibidi, Martinsried, Germany) placed on a 24-well glass-bottom plate coated with 0.3 mg mL^-1^ type I collagen. After 6 h of incubation, the insert was removed, and MDCK-4KO-EKARrEV-NLS cells were plated around the EGFRL-producer cells. Cells were supplemented with Marimastat 10 μM to inhibit the secretion of EGFRL before observation. After 2 h, cells were washed with PBS to remove the EGFRL-receiver cell aggregates on EGFRL-producer cells and the medium was replaced with Medium 199 supplemented with Marimastat 10 μM, 10% FBS, 100 units mL^-1^ penicillin, and 100 µg mL^-1^ streptomycin. After 166 h, the interface between EGFRL-producer cells and MDCK-4KO-EKARrEV-NLS cells was imaged, and m^D^cTMP (final 10 μM) was added into the medium for the EGFRL secretion. To determine the cell displacement, the Fiji TrackMate plugin was applied to the CFP fluorescence images for tracking each cell over 10 h after treatment with m^D^cTMP.

### Analysis of ERK activity and full width at half maximum of ERK activation in each receiver cell

Shedding of HBEGF or EREG and observation of ERK activity by SLIPT was done as described above. For the analysis of ERK activity in each receiver cell, the velocity of the radial ERK activity wave was analyzed using MATLAB as described above, and time-series data of the FRET/CFP ratio and distance from the center were obtained.

Time-series data of the FRET/CFP ratio for 10 cells from the center were plotted against time for HBEGF and EREG, respectively. For quantification of the FWHM of ERK activation, the time required for ERK activation to recover to the half-maximum value was calculated. The half-maximum was defined as the average of the ratio value before EGFRL secretion (basal) and that just after adding m^D^cTMP (maximum) in each receiver cell.

### Fluorescence imaging of the endocytic pathway

For co-immunostaining of EEA1 and Rab7, 1.0 x 10^2^ MDCK-4KO-EREG-ScNeo or MDCK-4KO-HBEGF-ScNeo cells expressing miRFP703-eDHFR(69K6)-cRaf were mixed with 8.0 x 10^4^ MDCK-4KO cells and seeded on a µ-Plate 96-Well Black (no. 89626; ibidi). Cells were supplemented with Marimastat 10 μM to inhibit the secretion of EGFRL before observation. One day after seeding, cells were imaged at a resolution of 0.20 μm/pixel under a spinning-disk confocal Marianas system based on the Zeiss Axio Observer Z1 inverted fluorescence microscope and CSU-W1 spinning disk, equipped with 405-, 445-, 488-, 515-, 561-, and 640-nm lasers, a 63×/1.4 NA oil immersion objective, an Evolve electron-multiplying charge-coupled device camera, and piezo-controlled z-step motor, all controlled by SlideBook 6 software (Intelligent Imaging Innovation, Denver, CO). Typically, a z-stack of 20 x–y confocal images was acquired at 0.4 µm steps. During observation, m^D^cTMP was added to 10 μM. After 75 min, cells were fixed in freshly prepared 4% PFA for 30 min, permeabilized in 0.1% Triton X-100 in calcium- and magnesium-free (CMF)-PBS/0.1% BSA for 10 min, and then incubated for 1 h at RT with anti-EEA1 mouse antibody (1:250) and anti-Rab7 rabbit antibody (1:100) in CMF-PBS/0.1% BSA. EEA1 antibody was detected using secondary anti-mouse antibody conjugated with AMCA (1:25). Rab7 antibody was detected using secondary anti-rabbit antibody conjugated with Cy5 (1:250).

For co-immunofluorescence of EEA1 and Rab7, 1.0 x 10^2^ MDCK-4KO-EREG-ScNeo or MDCK-4KO-HBEGF-ScNeo cells expressing miRFP703-eDHFR(69K6)-cRaf were mixed with 4.0 x 10^4^ MDCK-Erbock#5 cells expressing EGFR and seeded on a 96-well glass-bottom plate (Matsunami Glass) coated with 0.3 mg mL^-1^ type I collagen. Cells were supplemented with Marimastat 10 μM to inhibit the secretion of EGFRL before observation. One day after seeding, cells were imaged under the TCS SP8 microscope at a resolution of 0.10 μm/pixel using a 63x/1.40 NA objective, 569 nm excitation for 10% and 650 nm for 10% of a white light laser, and HyD SMD detectors for 579–640 nm and 660–750 nm for gain of 500%, respectively. Typically, a z-stack of 40–50 x–y confocal images was acquired in 0.3 µm steps. During observation, m^D^cTMP was added to 10 μM. After 60 min, cells were fixed in freshly prepared 4% PFA for 30 min, permeabilized in 0.1% Triton X-100 in calcium- and magnesium free (CMF)-PBS/0.1% BSA for 10 min, and then incubated for 1 h at RT with anti-EEA1 mouse antibody (1:100) and anti-Rab7 rabbit antibody (1:100) and anti-RFP rat antibody (1:500) and in CMF-PBS/0.1% BSA. EEA1 antibody was detected using Alexa 405-conjugated goat anti-mouse IgG (H+L) antibody (1:250). Rab7 antibody was detected using Alexa 647-conjugated goat anti-rabbit IgG (H+L) antibody (1:100). RFP antibody was detected using Alexa 546-conjugated goat anti-rat IgG (H+L) antibody (1:500).

### Measurement of the fraction of mScarlet-EGFRL colocalized with EEA1 or Rab7

To quantify the amount of mScarlet-EGFRL colocalized with EEA1 or Rab7, 3D images of cells were smoothed using a Gaussian filter with sigma of 1.0 pixels by Fiji. A segment mask was generated from background-subtracted images to select voxels detected through the 561-nm or 579-nm channel (total mScarlet-EGFRL). Additional segment masks were generated to include all voxels detected through the 640-nm or 660-nm channel (Mask-Rab7), and to include all voxels detected through the 405-nm channel (Mask-EEA1). For all masks, identical threshold parameters were used for experimental variables. A “colocalization” mask was then generated to select voxels overlapping between the total mScarlet-EGFRL mask and Mask-Rab7 or Mask-EEA1. The sum fluorescence intensity of the 561- or 579-nm channel in the colocalization mask was divided by the sum fluorescence intensity of the mScarlet-EGFRL in each FOV to calculate the fraction of total cellular mScarlet-EGFRL co-localized with EEA1 or Rab7.

### Generation of *Ereg* KO mice

The B6N Albino-*Ereg*^-/-^ Tg (hyBRET-ERK-NLS) pT2A-6011NLS (or simply, *Ereg*^-/-^ hyBRET-ERK-NLS) mice were developed using a CRISPR/Cas9 system targeting the murine *Ereg* gene (NC_000071.6). Three Alt-R™ CRISPR-Cas9 crRNAs (TableS2) were co-injected with Alt-R™ CRISPR-Cas9 tracrRNA and Alt-R™ S.p. Cas9 Nuclease V3 (Integrated DNA Technologies, Inc., Coralville, IA) into the cytoplasm of the fertilized eggs obtained from C57BL/6N hyBRET-ERK-NLS mice (Komatsu et al., 2018) as described previously (Sunagawa et al., 2016). The mice were deposited at the Laboratory Animal Resource Bank, National Institutes of Biomedical Innovation, Health and Nutrition (Osaka, Japan; hyBRET-ERK-NLS Ereg-/-, Resource no. 448).

### Genotyping of *Ereg* KO mice by quantitative PCR (qPCR)

Genomic DNAs from the fingers of wild-type (WT) and *Ereg* KO mice were prepared by isopropanol precipitation after lysis with Proteinase K (Nacalai Tesque), and then subjected to qPCR analysis for genotyping of the mice using the CFX Connect Real-Time PCR System (Bio-Rad Laboratories, Hercules, CA), TB Green® Premix Ex Taq™ II (Tli RNaseH Plus) (TAKARA bio) and primers for qPCR (TableS1). The absolute abundance of each target site was calculated using a standard curve obtained from WT genomic DNA. The amounts of target sites were normalized by the internal control, *Tbp* (Tsujino et al., 2013). When the amount of any of three target sites was less than 0.5% compared to the WT, the genotype was considered a KO.

### Time-lapse *in vivo* two-photon imaging of the wounded ear skin of mice

The *in vivo* imaging procedures were described previously (Hiratsuka et al., 2015; Komatsu et al., 2018). Briefly, 7- to 9-week-old mice were used for the *in vivo* imaging. The animal protocols were approved by the Animal Care and Use Committee of Kyoto University Graduate School of Medicine (approval nos. 22063, 23049). The experiments were carried out under the relevant regulations. Eighteen hours before the start of the imaging, mice were anesthetized with 1.5% isoflurane (Abbot Japan, Tokyo, Japan), the ear hair was removed, and a surgical scalpel was used to create epithelial wounds on the ear skin. Then, 2P excitation microscopy was performed with an FV1200MPE-IX83 inverted microscope equipped with a ×30/1.05 NA silicon oil-immersion objective lens (UPLSAPO 30XS; Evident), an InSight DeepSee Ultrafast laser (Spectra-Physics, Andover, MA), an IR cut filter (BA685RIF-3; Evident), two dichroic mirrors (DM505 and DM570; both from Evident), and two emission filters (BA460-500 for CFP and BA520-560 for YFP; both from Evident). The interval of the z-stack imaging was set at 1 μm. Kymographs depicting ERK activity were created via a customized MATLAB script. The Fiji TrackMate plugin was used to track each cell’s displacement over the course of 3 h based on CFP fluorescence images.

### Statistical analysis

All statistical analyses were carried out using Microsoft Excel software (Microsoft, Redmond, WA). Probability (p) values were determined by using the T.TEST function of Microsoft Excel with two-tailed distribution and two-sample unequal variance. The sample number for this calculation (n) is indicated in each figure legend.

## Supplemental figure legends

**Figure S1.**
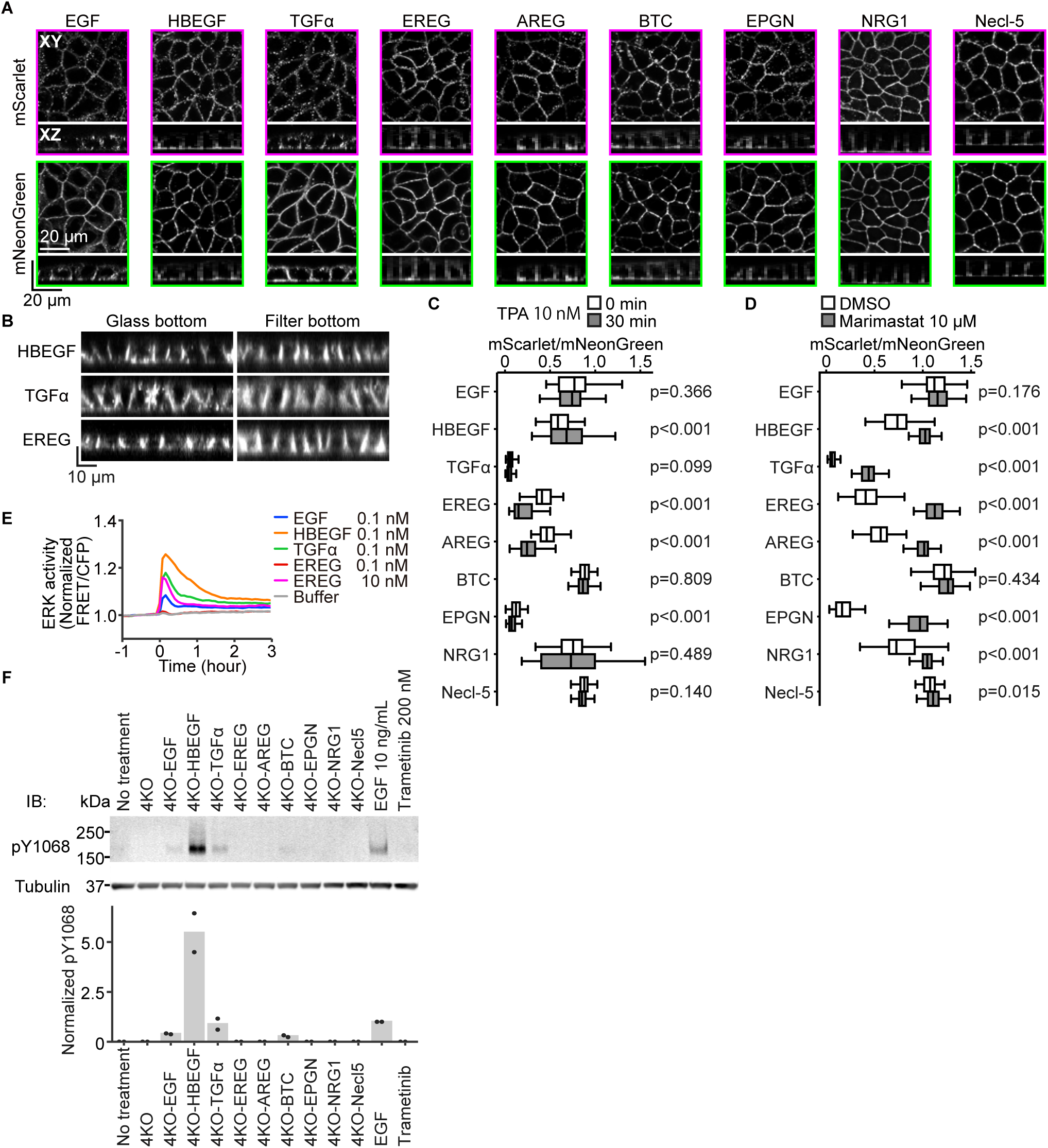
Dual color probes visualize subcellular localization, internalization, and shedding of EGFRLs. **(A)** Identical to Fig. 1C. Low-magnification XY images and XZ images are shown. **(B)** Confocal XZ projection mNeonGreen images of HBEGF-ScNeo, TGFα-ScNeo, and EREG-ScNeo-expressing MDCK cells plated either on the cover glass or permeable filter. **(C)** EGFRL-ScNeo-expressing cells were stimulated with 10 nM TPA for 30 min. The mScarlet/mNeonGreen ratio of the plasma membrane was quantified. Data were pooled from three independent experiments. From the images of three independent experiments, 84 cells under each condition were randomly selected for analysis. Data are shown in the box plot. **(D)** EGFRL-ScNeo-expressing cells were cultured in the presence of 0.1% DMSO or 10 μM Marimastat for one day before imaging. The mScarlet/mNeonGreen ratio of the plasma membrane was quantified. Data were pooled from three independent experiments. From the images of three independent experiments, 90 cells under each condition were randomly selected for analysis. Data are shown in the box plot. **(E)** MDCK-4KO-EKARrEV-NLS cells observed under a fluorescent microscope were stimulated with the 0.1 nM recombinant EGFRLs. The FRET/CFP ratio was normalized to the average values over the 60 min-observation period before stimulation. ERK activity (normalized FRET/CFP) was quantified and plotted as a function of time. Data were pooled from three independent experiments. n > 1000 cells for each condition. **(F)** Western blotting analysis of cell lysates of MDCK-4KO-EKARrEV-NLS cells stimulated with the supernatant of MDCK-4KO-EGFRL-ScNeo cells or 10 ng/mL EGF or 200 nM Trametinib. pY1068 was normalized with EGF condition. Each dot indicates an independent experiment. The bar graphs represents the means. Statistical significance was determined by unpaired two-tailed Welch’s t-test.

**Figure S2.**
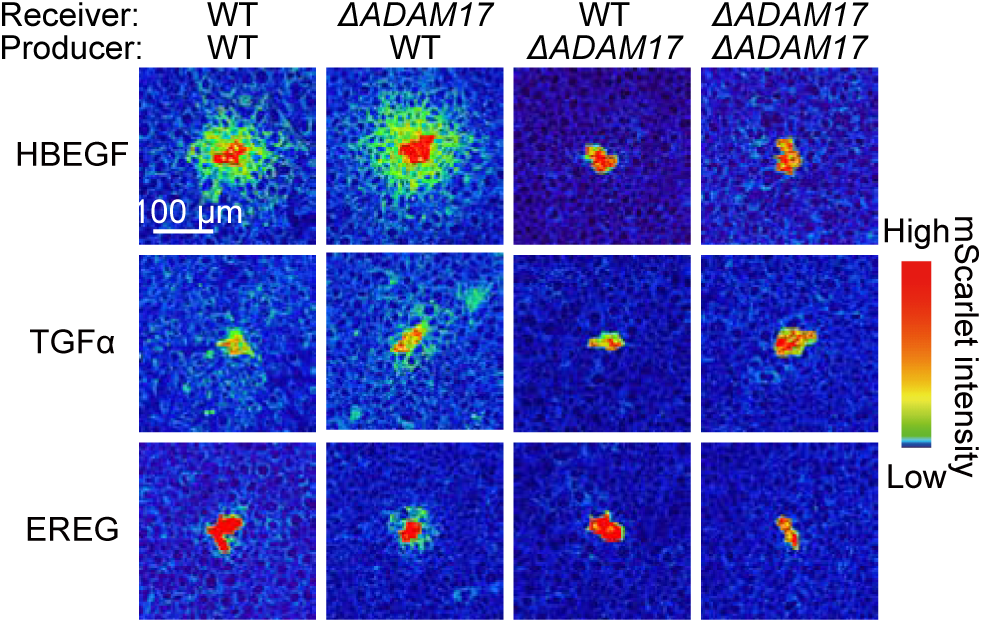
**ADAM17 is required only in the producer cells.** As in Fig. 2, the EGFRL-producer and -receiver cells were cultured at a ratio of 1:400. The producer and receiver cells were derived from either WT or ADAM17-knockout MDCK cells. Shown here are representative mScarlet confocal images acquired in a single plane at 24 h after seeding. The red-colored region indicates producer cells.

**Figure S3.**
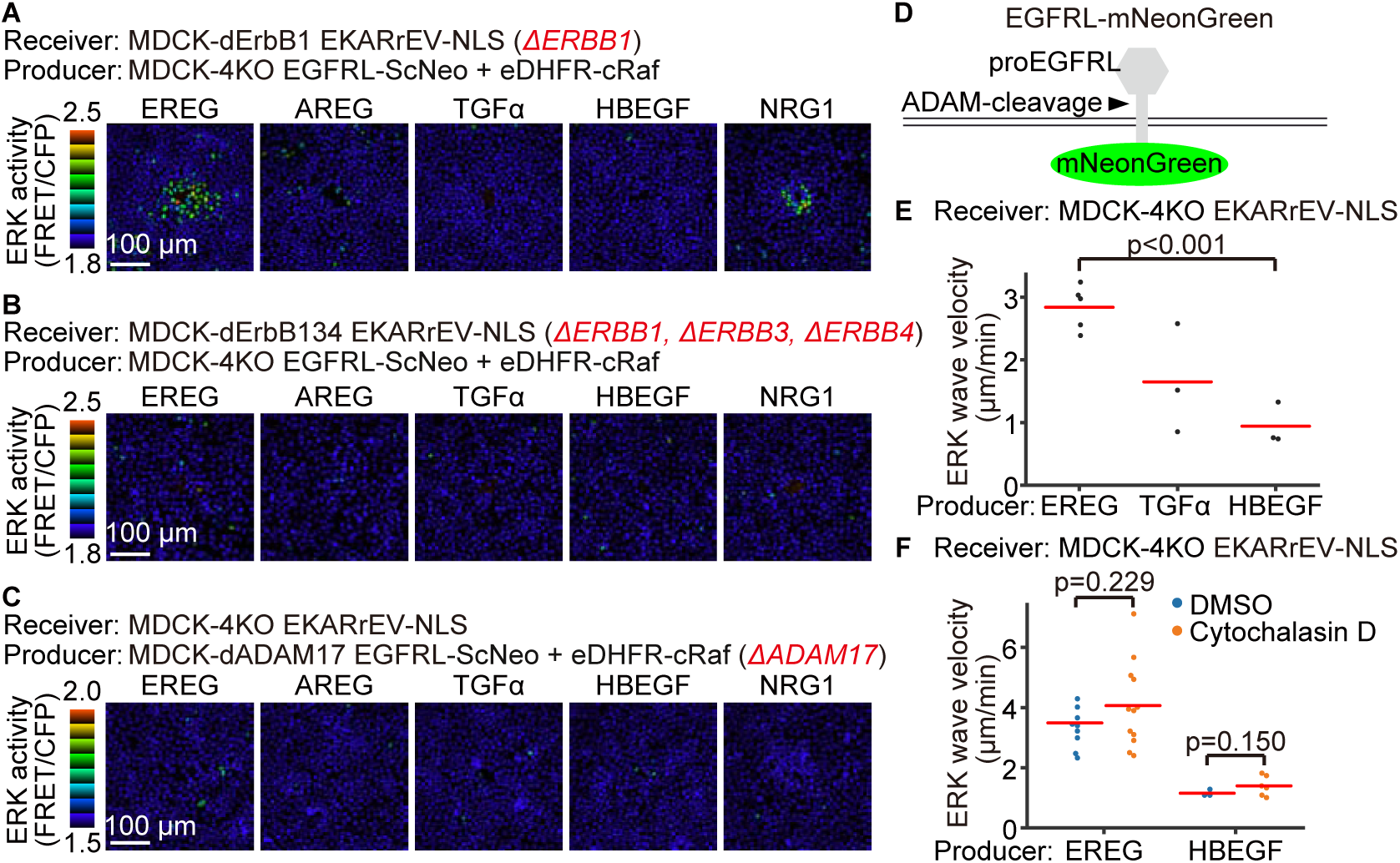
ADAM17 in the producer cells and ErbB receptors in the producer cells are required for ERK wave propagation by EGFRLs. **(A)** Representative images of ERK activity (FRET/CFP) in MDCK-dErbB1-EKARrEV-NLS receiver cells. The producer cells were located in the center of the image. Images were acquired 20 min after m^D^cTMP addition. **(B)** Representative images of ERK activity (FRET/CFP) in MDCK-dErbB134-EKARrEV-NLS receiver cells. The producer cells were located in the center of the image. Images were acquired 20 min after m^D^cTMP addition. **(C)** Representative images of ERK activity (FRET/CFP) in MDCK-4KO-EKARrEV-NLS receiver cells. ADAM17 of producer cells was deleted. The producer cells were located in the center of the image. Images were acquired 20 min after m^D^cTMP addition. **(D)** A schematic of EGFRL-mNeonGreen with mNeonGreen located in the cytoplasm. **(E)** The velocity of ERK wave propagated from producers expressing each ligand fused with C-terminal mNeonGreen. Each dot indicates a single producer cell population. The red bars represent the means. n = 5 (EREG), 3 (TGFα), and 3 (HBEGF) producer cell populations from a single experiment. **(F)** Maximum radius of ERK wave propagated from EREG- and HBEGF-producer cells to MDCK-4KO-EKARrEV-NLS receiver cells supplemented with 0.5% DMSO or 10 μM Cytochalasin D. Each dot indicates a single producer cell population. The red bars represent the means. n = 9 (EREG, DMSO), 12 (EREG, Cytochalasin D), 3 (HBEGF, DMSO), 6 (HBEGF, Cytochalasin D) producer cell populations from a single experiment. Statistical significance was determined by unpaired two-tailed Welch’s t-test.

**Figure S4.**
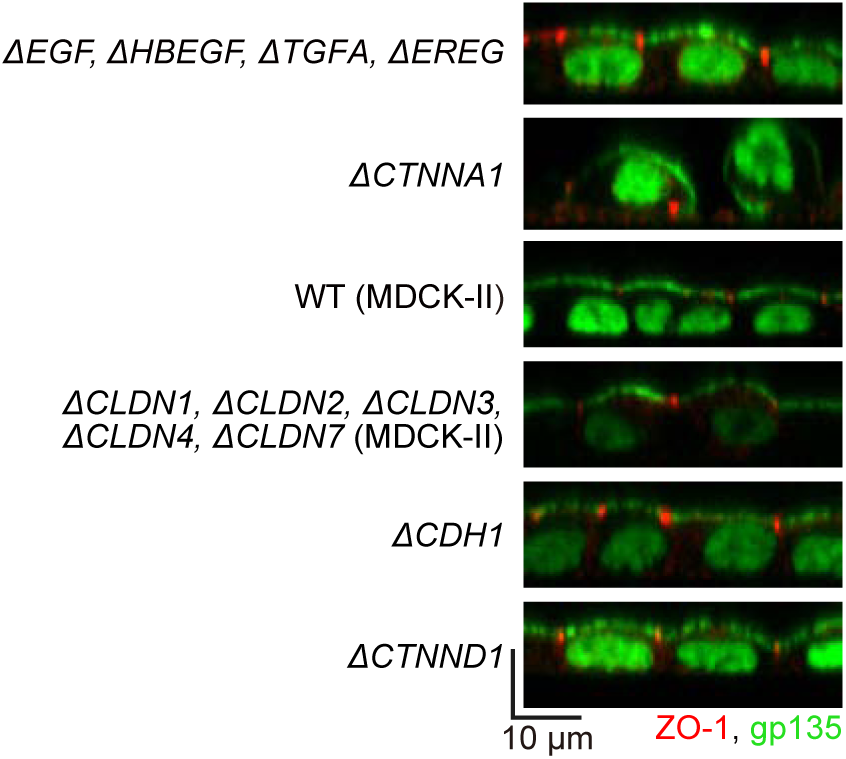
**Formation of a confluent epithelial layer in receiver cells** MDCK cells expressing ERK FRET biosensor were fixed and immunostained for ZO-1 (red) and gp135 (green). The YPet signal of the FRET biosensor in the nuclei is also depicted by the green channel. Confocal projections of XZ planes are shown.

**Figure S5.**
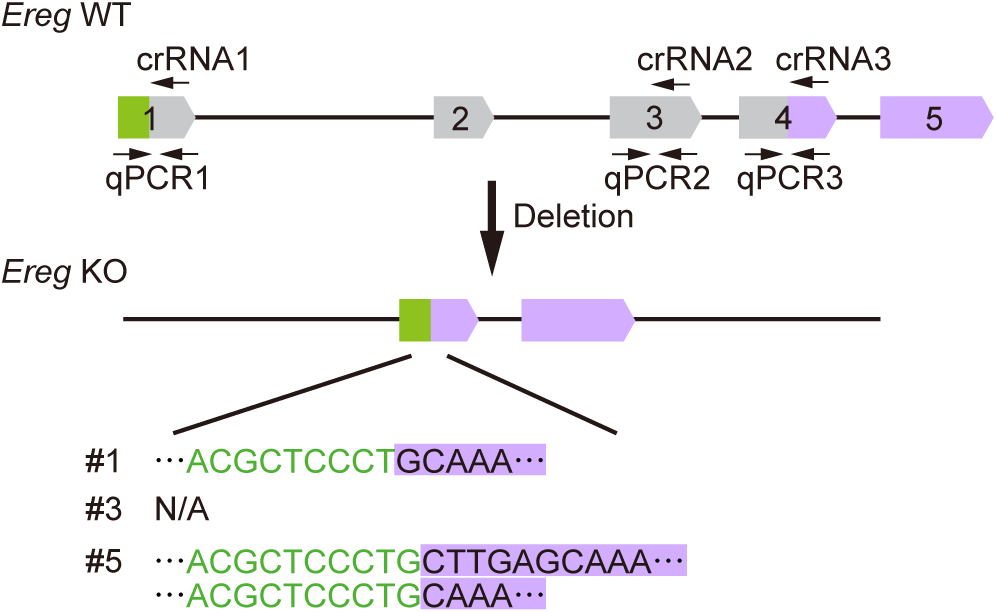
Generation of EREG-knockout mice. **(A)** Strategy for disrupting the mouse *Ereg* gene. Three crRNAs were used for CRISPR/Cas9-mediated mutagenesis. The positions of three pairs of qPCR primers for the genotyping are shown.

## Data availability

The data that support the findings of this study are available within the article and its Supplementary Information or from the corresponding author upon reasonable request.

**Video S1. Ectodomain shedding of EREG induced by TPA.**

The video image corresponds to the upper panel of Fig. 1I. The mScarlet/mNeonGreen ratio images of EREG-ScNeo-expressing MDCK cells are shown. Cells were stimulated with 10 nM TPA.

**Video S2. Inhibition of ectodomain shedding of EREG by ADAM17 inhibitor.**

The video image corresponds to the lower panel of Fig. 1I. mScarlet/mNeonGreen ratio images of EREG-ScNeo-expressing MDCK cells are shown. The ADAM17 inhibitor Marimastat was added to 10 μM.

**Video S3. Ectodomain shedding of AREG triggered by cRaf activation.**

Time-lapse movies of miRFP703 and mScarlet/mNeonGreen ratio images (Fig. 3C). The AREG-ScNeo probe was introduced into the miRFP703-eDHFR-cRaf cells. Cells were treated with 10 μM m^D^cTMP.

**Video S4. Propagation of ERK activation by EGFRLs.**

Time-lapse movies of FRET/CFP ratio images (Fig. 3F). The producer cells expressing EGFRL-ScNeo and eDHFR-cRaf were co-cultured with an excess of MDCK-4KO cells expressing EKARrEV-NLS. Each ligand producer is located at the center of the image. m^D^cTMP 10 μM was added at time 0.

**Video S5. Propagation of ERK activation by HBEGF in α-1-catenin KO MDCK cells.**

Time-lapse movies of FRET/CFP ratio images (Fig. 4D). The producer cells expressing EGFRL-ScNeo and eDHFR-cRaf were co-cultured with an excess of α-1-catenin KO MDCK cells expressing EKARrEV-NLS. Each ligand producer is located at the center of the image. m^D^cTMP 10 μM was added at time 0.

**Video S6. Cell movement is driven by HBEGF-but not EREG-induced ERK waves.**

Time-lapse movies of FRET/CFP ratio images (Fig. 5B). The producer cells expressing EGFRL-ScNeo and eDHFR-cRaf (left) were co-cultured with MDCK-4KO cells expressing EKARrEV-NLS. m^D^cTMP 10 μM was added at time 0.

**Video S7. Activity propagation of ERK, tyrosine kinases, and ROCK.**

Time-lapse movies of FRET/CFP ratio images (Fig. 5D). The producer cells expressing EGFRL-ScNeo and eDHFR-cRaf were co-cultured with an excess of MDCK-4KO receiver cells expressing EKARrEV-NLS, Picchu, or Eevee-ROCK. m^D^cTMP 10 μM was added at time 0.

**Video S8. ERK waves in WT and EREG deficient mice.**

Time-lapse movies of FRET/CFP ratio images (Fig. 7B). With hyBRET-ERK-NLS-expressing transgenic mice with a WT or an *Ereg^-/-^*background, a surgical scalpel was used to create epithelial wounds on the ear skin. Mice were observed under two-photon excitation microscopes. Three mice of each background were observed.

**Supplementary Table S1. Primers used in this study.**

**Supplementary Table S2. sgRNAs and crRNAs in this study.**

