## Supplementary Table S2 for "Low-affinity ligands of the epidermal growth factor receptor are long-range signal transmitters during collective cell migration of epithelial cells"

| Target | Cell | Sequence | Source | Additional notes |
| --- | --- | --- | --- | --- |
| EGF-KO | MDCK | 5’- CTTATCATTCTGTGGCCGGT -3’ | Lin et al., 2021 | sgRNA |
| HBEGF-KO | MDCK | 5’- CTCCCACCGAATCCACGGAC -3’ | Lin et al., 2021 | sgRNA |
| TGFα-KO | MDCK | 5’- GTCCCACTTCAACGACTGCC -3’ | Lin et al., 2021 | sgRNA |
| EREG-KO | MDCK | 5’- GATAGAAGACAACCCACGTG -3’ | Lin et al., 2021 | sgRNA |
| ErbB1-KO | MDCK | 5’- CCCAGGAGGCGGCACACGTG -3’ | Matsuda et al., 2023 | sgRNA |
| ErbB2-KO | MDCK | 5’- CAGAGGCTACGAATTGTGCG -3’ | Matsuda et al., 2023 | sgRNA |
| ErbB3-KO | MDCK | 5’- CCGACTCTTCAATGACAGCG -3’ | Matsuda et al., 2023 | sgRNA |
| ErbB4-KO | MDCK | 5’- ATTGATGTCGGCTCAGACTG -3’ | Matsuda et al., 2023 | sgRNA |
| ADAM17-KO | MDCK | 5’- TGGTGAAAGGCACTATAACA -3’ | Lin et al., 2021 | sgRNA |
| α-1-catenin-KO | MDCK | 5’- GTAGAAGATGTTCGAAAACA -3’ | Hino et al., 2022 | sgRNA |
| E-cadherin-KO | MDCK | 5’- CGGGGGCGCCGCCGTACCGA -3’ | This paper | sgRNA |
| p120-catenin-KO | MDCK | 5’- GGGCGTGACTTCCGCAAGAA -3’ | This paper | sgRNA |
| EREG-KO | B6 albino mouse | 5’- GCGTCAAGACCCAAGAGGCA -3’ | This paper | crRNA |
|  |  | 5’- CGTATTCTTTGCTCAAGGGT -3’ | This paper | crRNA |
|  |  | 5’- ATCTGCACTTGAGCCACACG -3’ | This paper | crRNA |

Table S2: Target sequences of sgRNAs or crRNAs for the gene knockout.
