## Supplementary Table S1 for "Low-affinity ligands of the epidermal growth factor receptor are long-range signal transmitters during collective cell migration of epithelial cells"

| Target | Cell | Sequence1 | Sequence2 | Source |
| --- | --- | --- | --- | --- |
| EGF-KO | MDCK | 5’- CTGAACGTCCGTTTCAGGCT -3’ | 5’- ACACTCCGAGAGAAATCAGGC -3’ | Lin et al., 2021 |
| HBEGF-KO | MDCK | 5’- GGGTGCAGAGGGGTCTGATG -3’ | 5’- GCAAGCTTGTCGGGTTTAGC -3’ | Lin et al., 2021 |
| TGFα-KO | MDCK | 5’- TGCAAAAGTTAAGGTGCGGC -3’ | 5’- TTTGAAGCAGGTGTCGCTCA -3’ | Lin et al., 2021 |
| EREG-KO | MDCK | 5’- CCTTAGGGGTCAAATGTTTGTTTAGTGGTG -3’ | 5’- GCTTTGTAAGTTTTGTAGGTTTGAGCTGCC -3’ | Lin et al., 2021 |
| ErbB1-KO | MDCK | 5’- TGCCTGGGTGTCTCTGTGAAGGGAAATCCT -3’ | 5’- CCCACCTGTCCACTTTCCGGCTGTCTTTAT -3’ | Matsuda et al., 2023 |
| ErbB2-KO | MDCK | 5’- GGTTTTGCCCAAGATCCCCA -3’ | 5’- CCACCTGCATCCATCCTGAA -3’ | Matsuda et al., 2023 |
| ErbB3-KO | MDCK | 5’- GGCCCTCAGGATACAGACTG -3’ | 5’- CAGCTGGCTACACAAACTCC -3’ | Matsuda et al., 2023 |
| ErbB4-KO | MDCK | 5’- TGTGTGGGTATCCTGGGACTTG -3’ | 5’- TAGGGCAGAACTCATCTGGCAC -3’ | Matsuda et al., 2023 |
| ADAM17-KO | MDCK | 5’- GTGGAGTGCAGTGACAGACA -3’ | 5’- TATCTTGCTGTCTGCCCTGC -3’ | Lin et al., 2021 |
| EREG-KO | B6 albino mouse | 5’- GGATGGAGACGCTCCCTGCC -3’ | 5’- GGGCGTCGAGGGCTTACCTA -3’ | This paper |
|  | B6 albino mouse | 5’- TCAACGCAACGTATTCTTTGCTCAAGG -3’ | 5’- GTGGGCTACACTGGTCTGCGA -3’ | This paper |
|  | B6 albino mouse | 5’- GATGGAAGACGATCCCCGTG -3’ | 5’- AGTAGCCGTCCATGTCAGAACTAC -3’ | This paper |
| Tbp (internal control) | B6 albino mouse | 5’- CCCCCTCTGCACTGAAATCA -3’ | 5’- GTAGCAGCACAGAGCAAGCAA -3’ | Tsujino et al., 2013 |

**Table S1**: Primers for validating the gene knockout.
